# Proximity determines donor candidacy during DNA double-stranded break homology directed repair

**DOI:** 10.1101/2025.02.10.637161

**Authors:** Charles D Yeh, Lilly van de Venn, Susanne Kreutzer, Xinhe Zheng, Naomi C. Cantos, Markus Schröder, Robin Hofmann, Felix E. Gerbaldo, Alexandra Clemens, Beeke Wienert, Christopher D. Richardson, Zacharias Kontarakis, Jacob E. Corn

**Affiliations:** ETH Zürich, Department of Biology, Institute of Molecular Health Sciences (IMHS); Zürich, 8093, Switzerland; ETH Zürich, Department of Biology, Genome Engineering and Measurement Lab (GEML); Zürich, 8093, Switzerland; University of California, Berkeley, Department of Molecular and Cell Biology (MCB); Berkeley, 94720, USA

## Abstract

DNA double-stranded breaks (DSBs) are especially toxic events that can be reversed by homology-directed repair (HDR), wherein information is copied from an intact template molecule. RAD51 mediates initial DSB/template pairing during homology search. A major challenge in understanding homology search in cells is the lack of tools to monitor this process. We developed **<u>RA</u>**D51 **<u>p</u>**roximity **<u>id</u>**entification **<u>seq</u>**uencing (RaPID-seq), a sensitive method that marks all candidate templates searched by RAD51. We find that HDR is hierarchical, such that DSB proximity determines template candidacy and subsequent recombination is unlocked by DSB/template homology. Sequences that lie outside the proximal window are not efficiently searched, even if identical in sequence. Our data reveal the invisible process of homology search and shed new light on fundamental mechanisms underlying genome editing.

## Main Text

Maintaining genome integrity is a vital function for living organisms and uncorrected DNA damage leads to malfunction and pathogenesis. DNA damage takes on a wide variety of forms which are reversed by an equally diverse number of repair pathways. Double-stranded breaks (DSBs) are among the most toxic DNA damage events. The two resulting DNA ends can potentially become uncoupled from one another, resulting in the loss of up to an entire chromosome arm (*1*).

The three major eukaryotic DSB repair pathways are non-homologous end-joining (NHEJ), microhomology-mediated end-joining (MMEJ), and homology-directed repair (HDR). While NHEJ and MMEJ are rapid, they can also introduce short insertion and deletion (indel) mutations. To ensure maximum genome integrity, HDR involves copying information from an intact template DNA donor, such as the sister chromatid formed during S phase. Alternatively, recombination between homologous chromosomes is possible, albeit with potential loss-of-heterozygosity (*2, 3*).

HDR requires extensive DSB end-processing and a laborious template-finding process called homology search. A filamentous homo-polymer of RAD51 is loaded onto processed single stranded DNA (ssDNA) on either side of the DSB. This nucleoprotein filament (NPF) then invades intact DNA duplexes to evaluate whether they are homologous to the DSB. There are two dominant models for homology search. Proximity-based models propose that homology search is mostly driven by the distance of candidate templates from the DSB. Homology-based models instead propose that the entire genome is available as a candidate template, with rapid evaluation and discarding of low homology sequences. These models have mostly emerged from cellular assays that measure stable binding of RAD51, cellular assays that track movement of RAD51 filaments, *in vitro* assays that measure RAD51 activity on small DNA fragments, or observation of the final sequences that emerge as end-products from the entire cascade of HDR (*3*–*6*). To measure the transient process of homology search in cells, we developed **<u>RA</u>**D51 **<u>p</u>**roximity **<u>id</u>**entification **<u>seq</u>**uencing (RaPID-seq).

Using RaPID-seq, we monitored double-stranded break repair (DSBR) homology search in human cells and found that only DNA in the immediate vicinity of DSBs was efficiently searched and recombined. Notably, the search profile is highly reproducible and unique for each genomic locus. Comparing homology search to chromatin conformation revealed that unperturbed cell chromatin conformation constrains search, including enabling search to distant regions. DSB-proximal endogenous DNA and high-concentration plasmids were efficiently searched, regardless of their homology to sequences at the break site. High homology was solely required as a second step to unlock final recombination after successful local search. Our results suggest that efficient HDR search is limited to the break proximal region. In the normal cell context, this constraint would favor recombination between the sister chromatids present during the S/G2 cell cycle phases and prevent aberrant recombination to distant genomic loci. However, exogenous DNA, such as those used in genomic editing contexts, can circumvent this restriction to serve as an efficient HDR donor template.

## Results

### RaPID-seq is a highly sensitive method for monitoring HDR homology search

RaPID-seq adapts the DamID DNA-protein proximity-labelling system to identify interactions between the HDR homology search filament and potential template DNAs. We fused RAD51 to the bacterial protein DNA adenine methyltransferase (*Dam*), which methylates double stranded DNA GATC motifs to Gm6ATC (*2, 7*). During HDR, this Dam-RAD51 (RaPID) fusion is incorporated into the homology search nucleoprotein ssDNA-RAD51 complex. As HDR search proceeds, each candidate genomic locus contacted by the NPF is irreversibly methylated, and these labeled molecules are identified by next generation sequencing DamID-seq (*8*). Combined with targeted deep sequencing (amplicon-NGS) of DSB sites, the terminal outcomes of DSBR can also be determined from the same cell population (**Fig. 1a**).

**Fig. 1.**
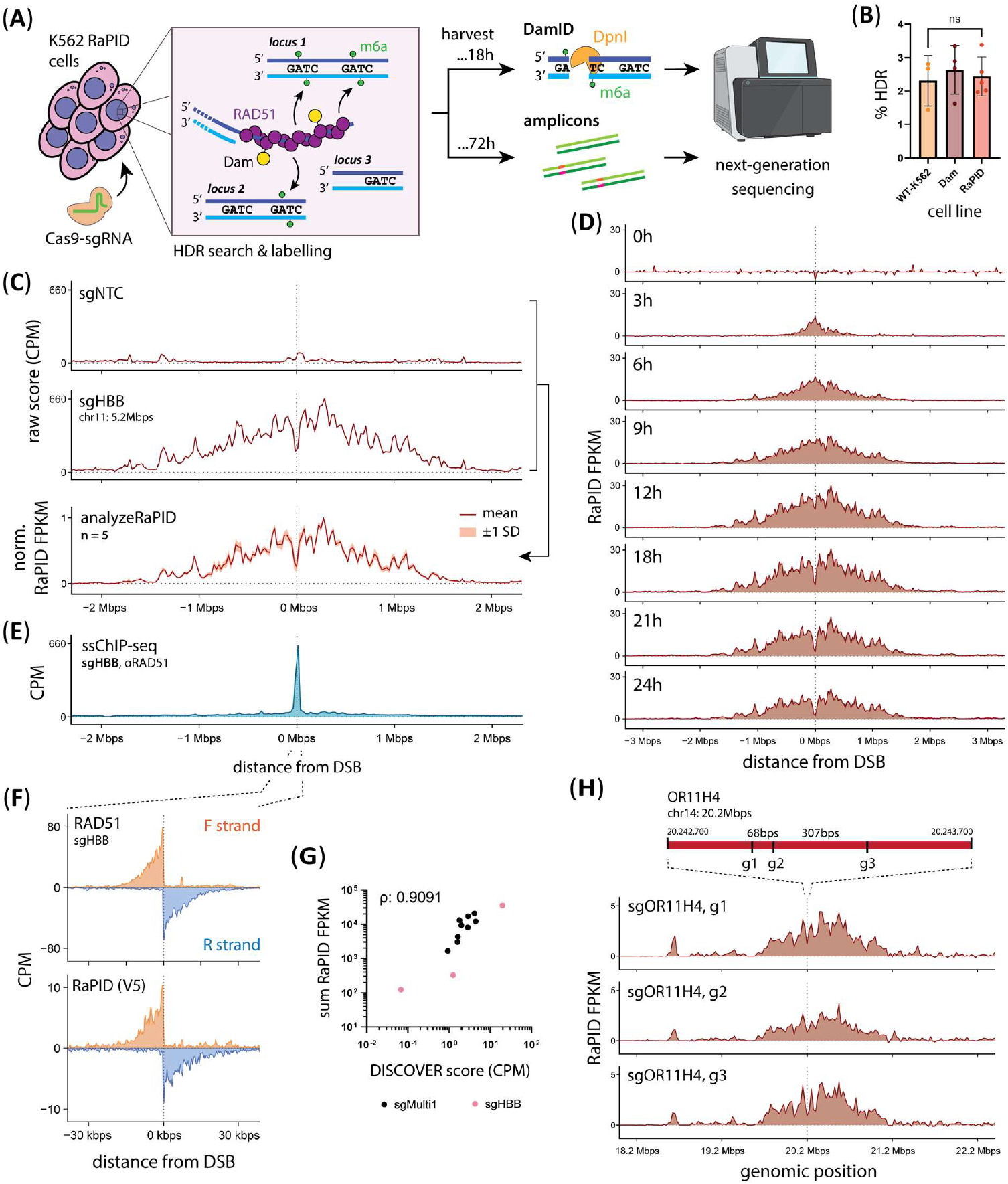
RaPID-seq enables highly sensitive and cumulative monitoring of HDR homology search *in situ/in vivo*. **(a)** Model for Dam-RAD51 (RaPID) labeling of probed loci during HDR search and workflow to resolve both HDR *candidacy* (DamID) and *choice* (amplicons) information. **(b)** Comparison of endogenous *HBB-HBD* recombination rates between cell lines. **(c)** Exemplar result of DamID-NGS at *HBB* ±DSB at the 18h post-Nucleofection timepoint; *analyzeRaPID* combines ±DSB data to enable highly sensitive and reproducible homology search measurements; band is ±1 SD. **(d)** RaPID-seq timecourse for a DSB at *HBB*. **(e)** RAD51-ssChIP shows homology search complex formation at the DSB±20kbps region. **(f)** Both RAD51 and RaPID are recruited to the DSB site in a strand-specific manner. **(g)** Correlation (Spearman’s ρ) between total RaPID-seq signal (DSB±2Mbps) and normalized DISCOVER score for *sgHBB* and *sgMulti1* CRISPR/Cas9 target sites; (p-value: 0.0001, ***). **(h)** Homology search profiles for different DSBs targeting *OR11H4*.

Both RAD51 overexpression and RAD51 fusions can negatively impact HDR (*9, 10*). We therefore implemented the destabilization-domain/Shield-1 (DD) and tetracycline-induction (TetOn) systems to carefully titrate RaPID expression such that it is present as a minority of the NPF (**Fig. S1a**) (*11*). To test RaPID-seq, we introduced a synchronized DSB using CRISPR/Cas9 protein and a well characterized *HBB*-targeting guide RNA (gRNA) as a ribonucleoprotein (RNP) complex in cells expressing either RaPID or Dam alone. A DSB at the targeted site in *HBB* results in endogenous recombination between *HBB* and *HBD*, which we measured by amplicon-NGS (*12, 13*). We found no difference in *HBB-HBD* recombination rates between WT-K562, Dam-K562, and RaPID-K562 cells, indicating that sub-stoichiometric introduction of the RAD51-Dam fusion is compatible with HDR (**Fig. 1b**). Using a Gm6ATC methylated Lambda-gDNA DamID spike-in control and a custom *analyzeRaPID* software pipeline (**Materials and Methods**), we developed a read depth-normalized RaPID score (RaPID FPKM) that enabled background subtraction and quantitative cross-comparison between samples.

RAD51-DamID marking occurred in both a DSB-dependent and RAD51-dependent manner (**Fig. 1c, S1bc**). Remarkably, potent methylation occurred in a ∼3Mbp window around the *HBB* DSB, with a short region lacking methylation immediately around the break site. We tested whether these were general properties of the RaPID-seq homology search signal using a multi-targeting guide RNA (sgMulti1) that targets the human genome 20 times across diverse genomic loci, as determined by DISCOVER-seq unbiased target identification (**Fig. S2a**) (*13*). Of these sites, we selected the 9 sites that had non-overlapping homology search profiles for further analysis. In all cases, the window of apparent homology search extended for megabases on either side of the DSB (**Fig. S1d**).

We found that RaPID-seq signal was highly reproducible across biological replicates at the same locus, suggesting that the observed homology search patterns are biologically informed rather than random variance (**Fig. 1c**). Gm6ATC is a persistent DNA mark, and we used RaPID-seq to measure the time-resolved cumulative homology search space. We found that homology search first occurred most intensively in a region immediately around the DSB (±150kbps) and expanded to further distances (±1.5Mbps). Homology search reached a maximum between 12-21h after a CRISPR/Cas9 DSB (**Fig. 1d**), consistent with prior reports of most HDR completing within 12-24h post-DSB induction (*14*). We observed a slight decrease in RaPID-seq signal by 24h, consistent with cells completing DSBR and re-entering the cell cycle, whereby replication would dilute the Gm6ATC mark.

Already by 6h post-DSB induction, we noticed a short ±3kbps unmethylated region at the DSB that grew in length at later time points, eventually extending to ±20kbps (**Fig. 1d**). Since DamID can only monitor double-stranded DNA, we postulated that the unlabeled region might represent resected single stranded DNA directly loaded with RAD51. We examined direct RAD51 recruitment using strand-specific chromatin immunoprecipitation (ssChIP) with antibodies for a V5 epitope tag on the Dam-fused RAD51 construct and for RAD51 itself (detects both endogenous RAD51 and Dam-fused RAD51). RAD51 and V5 were equally recruited to the target site in a DSB-dependent manner and on opposite strands around the DSB, consistent with known mechanisms of end processing (**Fig. 1e-f; S1e-f**). Closely examining the RAD51 ChIP-seq data, we observed evidence of a peak shoulder that matched the extents of the RaPID-seq search window but was more than 20-fold weaker (**Fig. S1e**). Notably, the main peak of RAD51 recruitment was perfectly anti-correlated with the unlabeled region of RaPID-seq signal.

Cumulatively, our results established RaPID-seq as a highly sensitive and robust method for monitoring RAD51-mediated homology search in cells.

### Homology search is constrained by DSB proximity and is sequence-independent

Having established RaPID-seq, we examined the roles of DNA proximity and sequence in homology search. While DSB-template sequence homology plays an important role during strand invasion and synapsis, these processes are downstream of homology search (*3*).

Strikingly, homology search from the *HBB* (chr11: 5.2Mbps) targeted DSB was solely restricted to the cut loci with no additional marking throughout the genome (**Fig. 2, S2b**). Even when multiple DSBs were introduced using a multi-targeting gRNA (sgMulti1), homology search was solely located around DSB sites as identified by DISCOVER-seq (**Fig. S2a, S3**). As expected, homology search signal strongly correlated with DSB processing by the MRN complex, a requisite step for NPF formation (**Fig. 1g**) (*13*). Despite extensive manual and computational searching, we found no evidence of RaPID-seq signal above background at any location lacking a DSB.

**Fig. 2.**
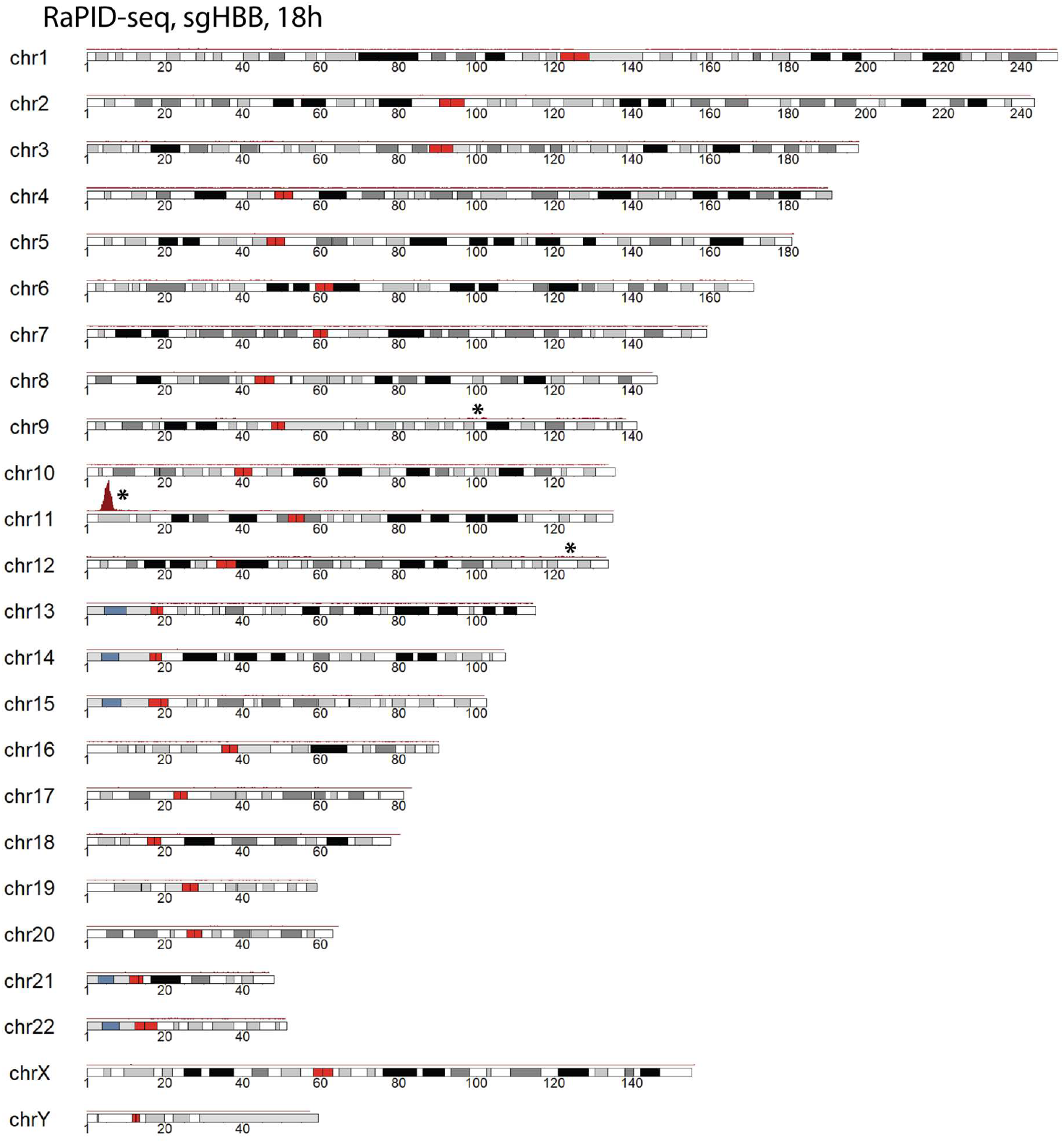
Genome-wide view of homology search profiles for CRISPR/Cas9 targeting with *sgHBB*. DSB target sites (*) were identified using DISCOVER-seq.

To assess if homology search profiles were affected by sequence at the DSB ends, we performed RaPID-seq for three different sites located within a kilobase of one another at the olfactory receptor (OR) gene *OR11H4*. Each of these sites has a distinct DSB end sequence, which could potentially alter a sequence-driven model of homology search (**Fig. S1g**). However, all three sites produced strikingly similar homology search profiles that were driven by proximity (**Fig. 1h**).

### Homology search and recombination success is constrained by pre-DSB chromatin conformation

Using RaPID-seq, we observed that homology search occurs proximal to the DSB with highly reproducible local profiles (**Fig. 1c, h**). As genomes are not 1-dimensional, we wondered if local chromatin conformation might influence homology search. To test this, we extracted one-to-all 4C “slices” (i.e., “virtual 4C”) from whole-genome all-to-all chromatin conformation (HiC) data for wild-type K562 cells at sites targeted by the several different *HBB* sgRNA, the sgMulti1 multi-targeting gRNA, OR-targeting gRNAs, and the commonly-used VEGFA-targeting gRNA (*15, 16*).

A naïve, 1-dimensional model of DNA proximity predicts a smooth power decay interaction frequency with the bulk of contact events occurring within 1Mbps of the DSB (**Fig. 3**, dashed line). However, virtual 4C exhibits substantial site-specific contact frequency variation that is driven by 3-dimensional DNA configurations (**Fig. 3, S4**). We found that the overall window of homology search occurred within DSB±1.5Mbps for all sites, as predicted by the naïve model. However, homology search often locally deviated from the naïve model. Local variation in search instead better matched chromatin contact frequency. Sites with especially high or low contact frequency had corresponding enhancements or reductions in homology search (**Fig. 3a**). This effect was especially dramatic at greater distances (>2Mbps), where the naïve model predicts minimal interaction. Homology search at these distances was matched by high distal chromatin contact frequency (**Fig. 3b**).

**Fig. 3.**
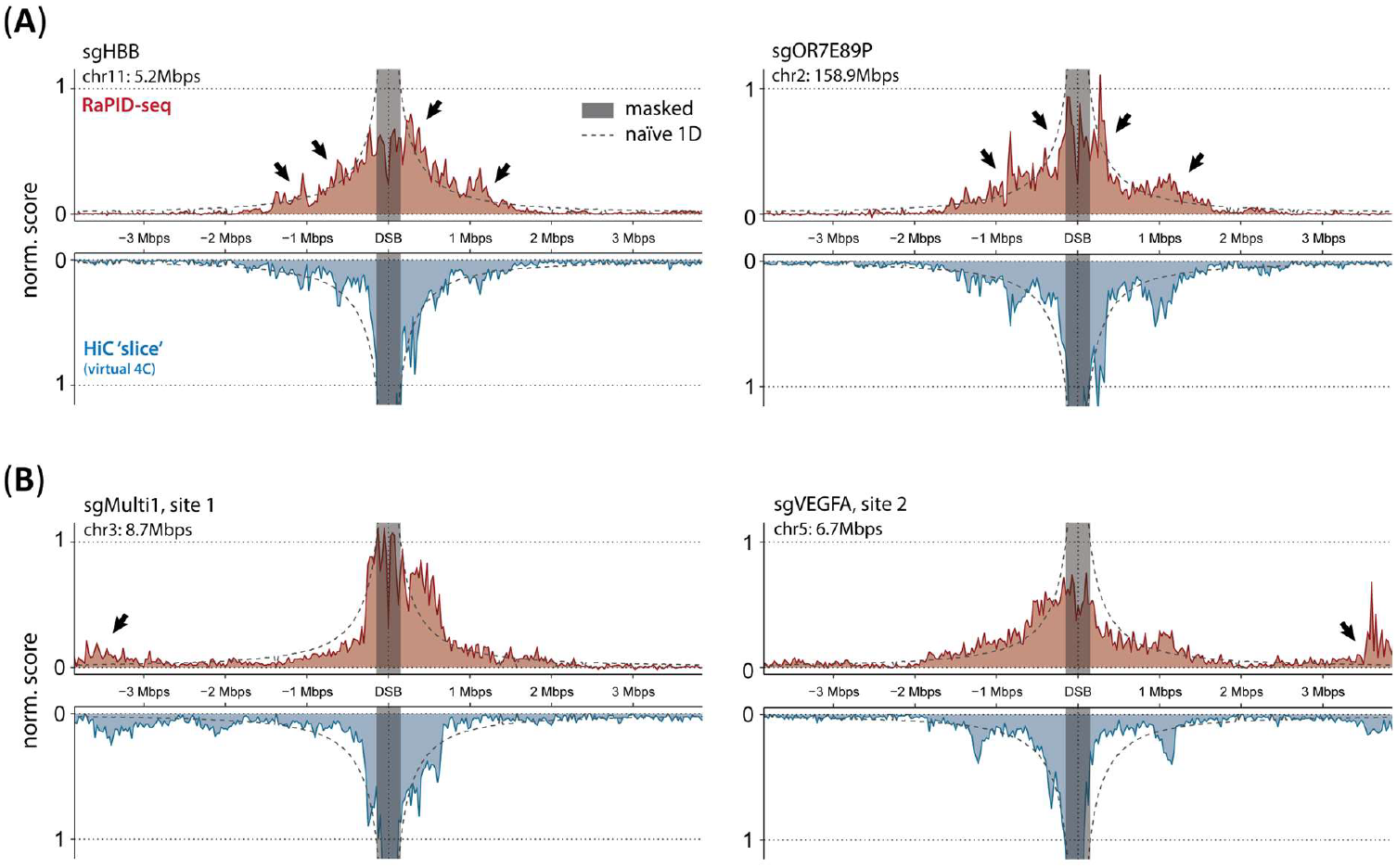
Comparison between unperturbed K562 cell chromatin conformation (HiC) and DSBR homology search (RaPID-seq) profiles. Dotted line curve: expected search profile for a naïve linear distance model of DNA interaction; masked region is DSB±150kbps. Deviation from the naïve model matches (arrows) in both chromatin conformation and homology search profiles at both **(a)** nearby distances (<2Mbps) and **(b)** at farther (>2Mbps) distances.

Importantly, the virtual 4C chromatin conformation data is from unperturbed cells, whereas RaPID-seq derives from a DSB event. Large-scale chromatin remodeling occurs in response to DSBs, including recruitment of DSBR proteins and changes in histone modifications (*17*). Despite these changes, we observed a striking similarity between pre-DSB chromatin conformation and homology search profiles. Our results therefore suggested that RAD51 search precedes chromatin remodeling during DSBR, such that pre-existing chromatin interactions have a strong influence on the template candidates during homology search and HDR.

### Proximity drives homology search to enable downstream recombination

To explicitly test the roles of proximity and homology in search, we used DSBs within human OR genes as a model. The OR genes are a collection of 874 related genes and pseudogenes of varying homology distributed across several chromosomes (*18*). A wide variety of templates of various homologies and distances are therefore available as templates for HDR when a DSB is formed in a given OR. We exploited this genetic and physical distance to ask whether high genomic homology or physical proximity dominates during homology search.

We focused on two distinct OR genes that recombine with a nearby OR gene after a CRISPR-Cas9 induced DSB, based on identifiable sequence signatures of the donor OR incorporated into the DSB-targeted OR: *OR7E39P* on chromosome 7 recombines with nearby *OR7E59P* and *OR7E89P* on chromosome 2 recombines with nearby *OR7E90P* (**Fig. S5b-c**).

RaPID-seq after a DSB at *OR7E39P* revealed a characteristic megabase-scale window of search around the break (**Fig. 4a, b left**), but no labeling or recombination with any OR genes elsewhere in the genome (**Fig. 4b right, S5b**). This is despite several distal OR genes having similar homology to *OR7E39P* as *OR7E59P* (**Fig. 4b right)**. OR genes located within the search window but having low homology to *OR7E39P* exhibited substantial labeling but were not recombined to the *OR7E39P* locus. RaPID-seq after a CRISPR-Cas9 DSB at *OR7E89P* yielded similar results. We observed a megabase-scale labeling window in *cis* to *OR7E89P*, with no evidence of search or recombination with highly homologous ORs outside this window on the same or distinct chromosomes (**Fig. 4c-d**). ORs with low homology but located within the *OR7E89P* search window were not recombined, and only *OR7E90P* was used as a template (**Fig. S5c**). In sum, targeted DSBs yielded RAD51 search in a 2-3Mbps local window, and only homologous sequences within this window recombined with the DSB site.

**Fig. 4.**
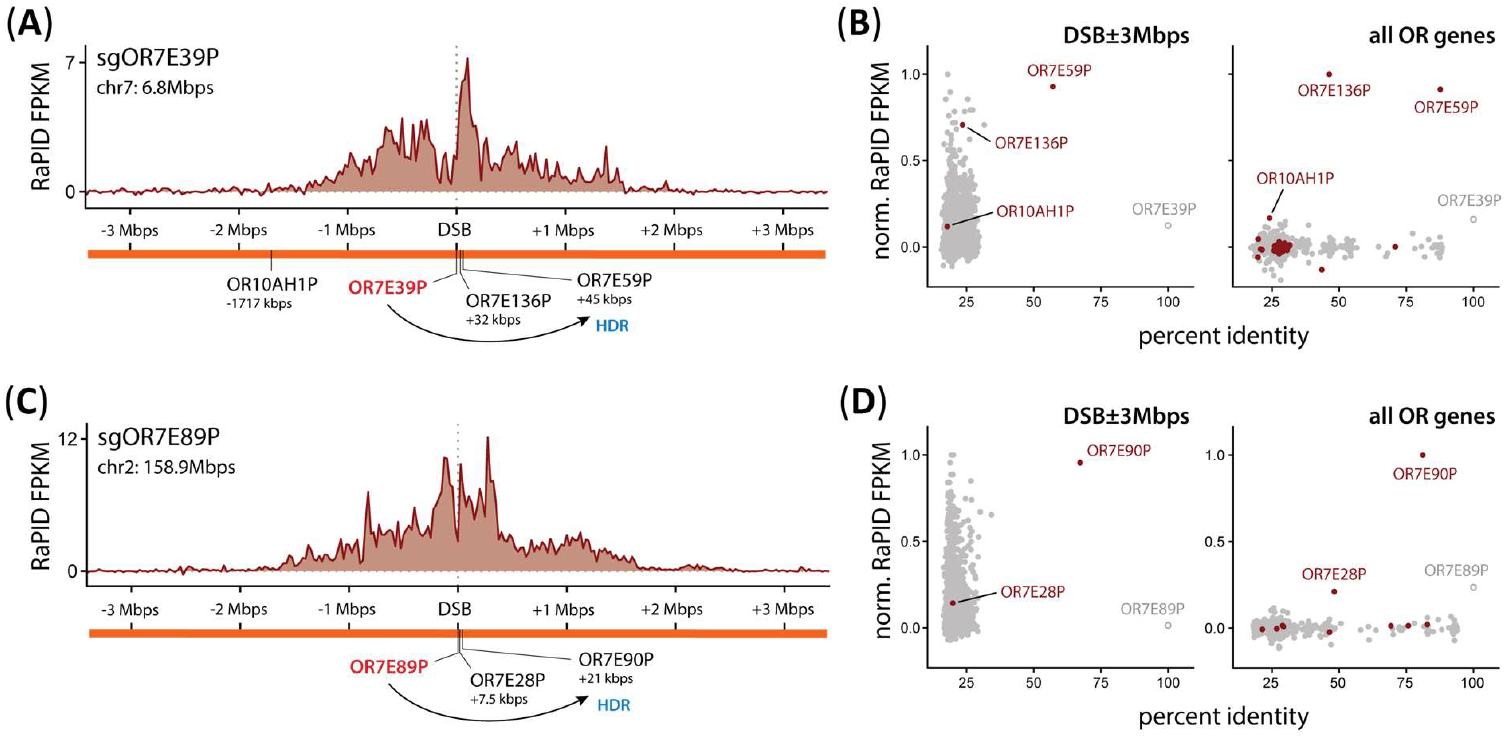
Endogenous HDR only occurs between the DSB and genomic loci that are both proximal and highly homologous. **(a)** RaPID-seq homology search profiles for CRISPR/Cas9 targeting at *OR7E39P*. **(b)** Comparison between normalized RaPID-seq score and sequence percent identity between the DSB at *OR7E39P* and local sequences within DSB±3Mbps region (left) and for all human OR genes (right). Highlighted points correspond to OR genes located on the same chromosome as the DSB. **(c, d)** Same as in **(a, b)** for CRISPR/Cas9 targeting at *OR7E89P*.

Our results so far suggested a local proximity-based model of homology search that determines donor candidacy, followed by unlocking of successful recombination for searched candidates with sufficient sequence homology. To test this model, we developed an orthogonal, one-to-all genome-wide recombination assay based on recombination of a DSB in BFP to a GFP donor template (*19*). The general goal of the assay was to determine all the locations in the genome from which equally homologous donor templates could recombine with a DSB.

We used low multiplicity of infection (MOI) lentiviral transduction to establish a monoclonal K562 cell line with a single copy integration of BFP under the control of a strong EF1α promoter, whose integration site (chr19:1Mbps) we mapped using a modified GUIDE-seq approach (*20*). We then further transduced this BFP clonal line with low MOI lentivirus bearing a non-expressed, promoterless GFP donor template to generate a polyclonal GFP donor pool having an estimated 480,000 different donor integration sites. Finally, we introduced a single CRISPR-Cas9 DSB at the expressed BFP and used fluorescence activated cell sorting to isolate cells where GFP recombined with the single copy of BFP (**Fig. 5ab, Table S1**). We used unique molecular identifiers to quantitatively map the locations of all GFP donors in this pool.

**Fig. 5.**
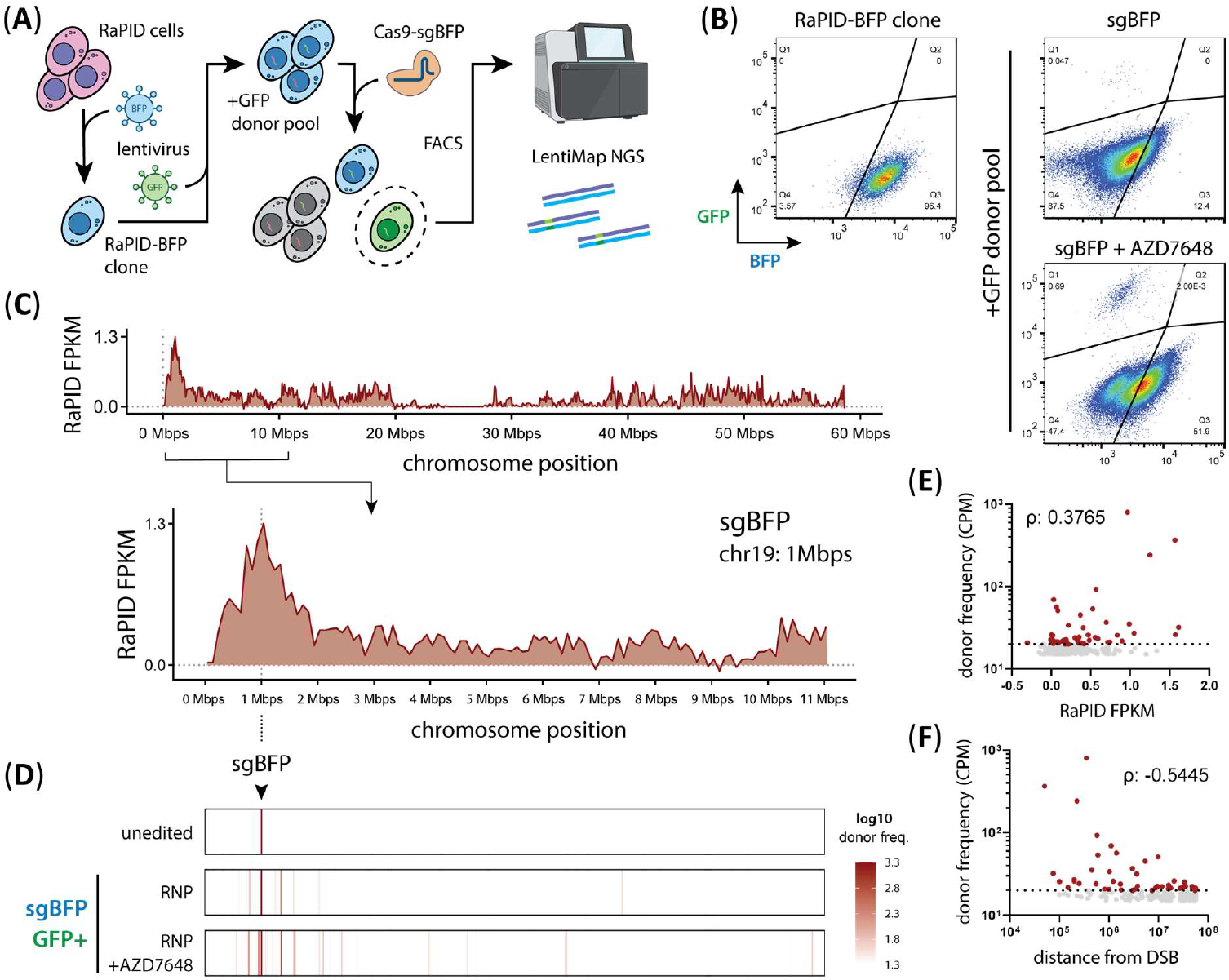
Unbiased genome-wide *in situ*/*in vivo* one-to-all HDR recombination analysis by *BFP* to *GFP* gene conversion. **(a)** Workflow for generating RaPID-BFP cell pools having random genome-wide *GFP* donor sequences by lentiviral transduction. DSBR at *BFP* results in either indel (non-fluorescent, gray) or HDR (green fluorescent) outcomes. **(b)** Flow cytometry for BFP and GFP. CRISPR/Cas9 targeting of *BFP* causes rare conversions to *GFP*, which are further enriched by DNA-PKcs inhibition (AZD7648). **(c)** RaPID-seq at *BFP* locus (chr19: 1Mbps). **(d)** Results of LentiMap assay for quantitative mapping of lentivirus insertion sites. Counts correspond to normalized abundance (CPM) of cells having a lentivirus insertion at the indicated locus. **(e)** Correlation (Spearman’s ρ) for sites passing threshold (highlighted points; donor frequency ≥ 20 CPM) between RaPID-seq score and genome-wide *BFP* to *GFP* recombination efficiency for AZD7648 condition (p-value: 0.0077, **) or **(f)** linear genomic distance for *GFP* donor loci (p-value: <0.0001, ****) for all sites on chromosome 19.

Using RaPID-seq, we found homology search from the single copy BFP-targeted DSB was concentrated in a megabase-scale DSB-proximal region, albeit with reduced signal relative to multiallelic endogenous genes such as *HBB* and the ORs (**Fig. 5c**). Targeting the same genomic region in parental non-BFP cells yielded a similar homology search pattern (**Fig. S6a**), indicating that the search window was independent of the BFP integration, consistent with our data on global chromatin conformation. Strikingly, GFP insertion sites within the RaPID-seq search window made up the vast majority of recombination events with *BFP*. This was despite hundreds of thousands of other equally homologous GFP donors being distributed throughout the genome across the unsorted cell pool (**Fig. S6b**).

We used inhibition of DNA-PKcs with AZD7648 to increase HDR levels and found that overall recombination rates increased 14-fold (*21*). However, this increase in HDR still predominantly stemmed from GFP donors located within the local search window (**Fig. 5b, d**). We also observed much rarer recombination between BFP and GFP donors mapped to chr9: 136.9Mbps and chr12: 48.7Mbps, which may suggest that genome-wide search can occur at very low frequencies in human cells (**Table S1**). Alternatively, K562 cell chromosomal rearrangements may have migrated distal genomic loci into the proximity of BFP (*22*). Recombination was in proportion to HDR search intensity (**Fig. 5d-e**), with increasing genomic distance between the BFP and GFP leading to a concomitant decrease in recombination rates (**Fig. 5f**).

Overall, our results so far supported a model wherein proximity to a DSB is the primary determinant of homology search, with homology of the DSB end to candidate templates having little effect on the searched region of the genome. Homology is instead required for downstream unlocking of successful recombination after proximal search.

### Exogenous DNA templates compete with endogenous genome search

During genome editing, HDR is used as a tool to enable highly precise genomic changes without unintended modifications. Typically, an exogenous DNA template encoding a desired genetic change is introduced to cells at the same time as a genome editing nuclease. The cellular HDR machinery then copies information from the donor into the genome editing site (*23*).

While our endogenous HDR experiments indicated that homology search is locally constrained, exogenous DNA templates such as ssDNAs or plasmids might bypass this requirement by nature of their abundance and mobility. To determine homology search dynamics in the context of exogenous DNA, we performed RaPID-seq in the presence of plasmid donors with varying homology to a CRISPR-Cas9 DSB targeted to the *HBB* gene (**Fig. 6a-b**). Reads aligning to the plasmid donor templates or the genome were computationally separated and analyzed to determine their search profiles. We performed these experiments in the context of DNA donor templates with varying homology to the DSB, ranging from 99.9% homology (only a PAM mutation, to prevent cutting of the donor by Cas9) to 0% homology.

**Fig. 6.**
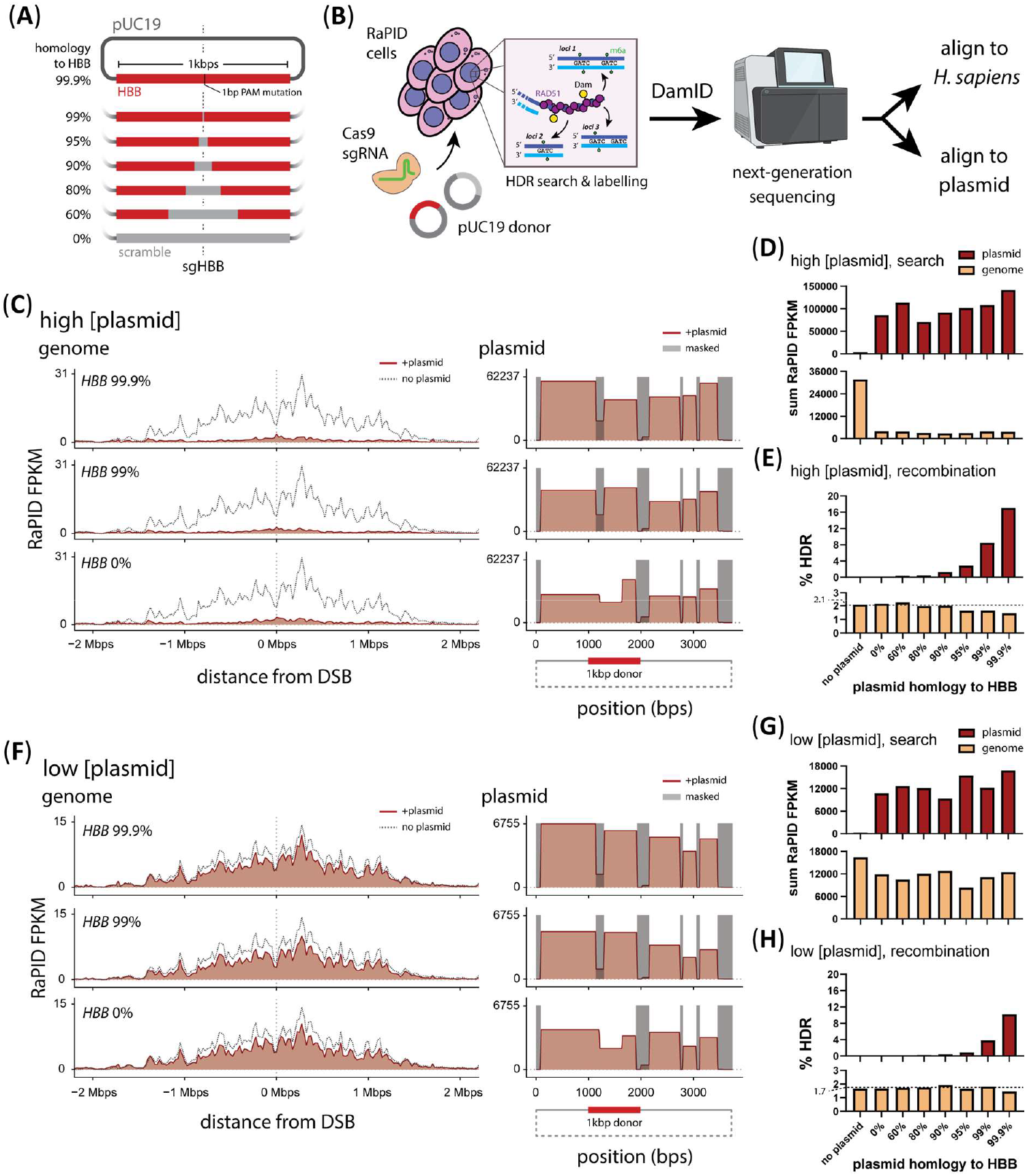
Exogenous DNA competes with endogenous DNA during DSBR homology search. **(a)** Design of plasmid DNA donors with varying homology to the CRISPR/Cas9 targeted *HBB* gene. Total plasmid size is 3.6kbps. **(b)** RaPID-seq workflow in cells treated with exogenous plasmid DNA. Homology search profiles for both the endogenous loci (*HBB*) and exogenous plasmid DNA in cells treated with **(c)** high (100ng/μL) donor plasmid concentrations; dotted line: homology search profile in absence of exogenous plasmid for comparison; non-amplifiable regions (<200bps) on plasmid are masked from analysis. **(d)** Summary of total RaPID-seq labelling for the endogenous loci region (DSB±2Mbps) and entire plasmid. **(e)** Summary of HDR rates for endogenous *HBB-HBD* and exogenous *HBB*-plasmid recombination. **(f-h)** Same as **(c-e)** in cells treated low (12.5ng/μL) donor plasmid concentrations.

We were surprised to find that all donor templates, regardless of homology, were efficiently searched by RAD51, as measured by potent methylation of the entire relatively small plasmid (**Fig. 6c, S7**). Plasmid search was dependent upon a DSB at the endogenous genomic locus, indicating that it stemmed from RAD51 loaded on the genome and not breakage of the plasmid (**Fig. S7a**). Plasmid search was only minimally affected by donor homology, and even completely non-homologous plasmids were potently searched at a frequency that was relatively independent of homology (**Fig. 6d, top**). Examining the genome where the DSB was located, we were surprised to find that exogenous plasmids strongly suppressed RAD51 homology search of the surrounding genomic window, including when using non-homologous plasmids (**Fig. 6c-d, S7b**). Despite homology-independent search of exogenous donors, we found via amplicon-NGS of the *HBB* locus that homology played a deciding role in their usage for HDR outcomes (**Fig. 6e**).

We hypothesized that the much higher concentration and mobility of plasmids relative to an endogenous locus might “distract” RAD51 from genomic search. To test this, we repeated our experiments with an 8-fold lower concentration of plasmid (**Fig. 6f, S7c**). In these conditions, the exogenous plasmids were still efficiently searched by RAD51, albeit with a signal approximately 10-fold lower than at higher concentrations (**Fig. 6g**). Plasmid search still occurred independent of plasmid homology. At these lower plasmid concentrations, we observed that genomic search was now mostly rescued. We observed robust homology search around the site of the DSB, though quantitatively lower than in the absence of an exogenous donor. As before, genomic search was reduced approximately equally, regardless of the exogenous donor homology (**Fig. 6g, S7c**). Likewise, high homology was required for observable HDR sequence outcomes (**Fig. 6h**).

The ability of high plasmid concentrations to diminish genomic marking suggests that RAD51 search might be limiting in cases where donor availability is very high, such as when supplying an exogenous donor during genome editing. These data emphasize the model in which RAD51 performs a greedy homology-independent search for appropriate donors that are physically located near the DSB, either by virtue of being at the *cis* locus or due to high concentration or mobility. Searched donors are then used in a homology-dependent manner.

## Discussion

We developed RaPID-seq to directly monitor the otherwise invisible process of RAD51 homology search during DSB-induced homology directed repair. Our data in human cells support a hierarchical model wherein HDR donor candidacy is determined by proximity, thereby limiting HDR donor choice. The homology-dependent usage of donors to yield a sequence-observable outcome is pre-constrained by this search space (**Fig. S8**).

We observed a striking similarity between unperturbed cell HiC data and DSB-induced RaPID-seq results. This suggests that HDR donor candidates must be proximal to the DSB before the DNA damage takes place. While DSBs can induce genomic reorganization, our data imply that reorganization does not dramatically expand homology search in human cells (*17, 24*). Genome reorganization may therefore be important for a downstream step of HDR. Notably, the sister chromatid is a perfectly matched homology donor, and the sisters are tethered together in S/G2 (where HDR is most active). Sister tethering dramatically increases the local concentration of a matched genomic locus, potentially pre-organizing the genome for efficient search of a sister around an equivalent region to the break and simultaneously disfavoring potentially damaging search of homologous distal locations such as repetitive elements (*3, 25*). While our system of CRISPR-Cas9 induced DSBs paired with RaPID-seq cannot separately resolve sister chromatid search, this would be an exciting future direction.

Despite sister chromatid tethering, exogenous DNA donors, such as plasmids used in genome editing experiments, can overcome this proximity constraint to efficiently compete with endogenous homology search. Potentially, this effect occurs by virtue of relatively high exogenous donor concentration and mobility (**Fig. S8**). Our observed concentration dependence of exogenous donor homology search is consistent with prior reports that increasing nuclear delivery of donor templates increases HDR (*26, 27*). Given that the presence of even non-homologous exogenous DNA donors affects homology search, other forms of DSB repair may also be affected. These as-of-yet unknown secondary effects of exogenous DNA may explain some reports wherein inclusion of even non-homologous exogenous DNA material affects genome editing outcomes (*28*).

As homology search is highly constrained by chromatin organization, tissue-specific differences in chromatin organization may affect HDR search outcomes. Several studies have already identified changes in chromatin organization in differentiating stem cells, including between different chromosomes (*29*–*32*). Under a proximity-based model, these chromatin organization differences could cause different homology search profiles, including enabling recombination between distant loci (*33*–*35*). Potentially, proximity-induced recombination between otherwise distal loci could play a role in recombination events that drive tissue-specific tumors.

Our results imply that human cells strongly favor local DSBR homology search over genome wide search. This is in striking contrast to bacteria and budding yeast, including recent descriptions of RAD51 filaments that stretch across the entire yeast nucleus (*5, 36*). Both the nucleus and genome are dramatically larger in human cells than in yeast, and human cells have overall reduced HDR rates in comparison (*3*). This does not preclude mechanisms of increased DSB end mobility during HDR in human cells, such as recruitment of nuclear actin (*37*). It may be that such processes do not increase the search space of donor candidates and instead ensure efficient and maximal homology search within the local region. We also note that a proximity-based model for search does not entirely preclude reported infrequent recombination between distant genomic loci (*33*–*35*). These events may occur rarely due to infrequent random contact, but they are in the minority as compared to the dominant mode of proximity-driven search. Measuring such rare interactions would require an even more sensitive technique than RaPID-seq at very high sampling depths. Further studies of rare long-range donor search could help understand genomic rearrangements that drive the outgrowth of rare clones in oncogenesis and/or improve genome editing applications.

## Data and materials availability

Computational data processing and analysis code for RaPID-seq is available online (https://github.com/yehcd/rapid-tools). Next-generation sequencing data is available at the Sequence Read Archive (SRA) repository under SUB15025610. Plasmids used in this study are available at Addgene (https://www.addgene.org/jacob_corn/).

## Notes

https://github.com/yehcd/rapid-tools

### Competing Interest Statement

JEC is a co-founder and scientific advisory board (SAB) member of Spotlight Therapeutics and Serac Biosciences and an SAB member of Mission Therapeutics, Relation Therapeutics, Hornet Bio, Kano Therapeutics and the Joint AstraZeneca CRUK Functional Genomics Centre. The laboratory of JEC has funded collaborations with Allogene, Cimeio, CSL Behring and Serac. None of these collaborations are related to this paper.

