## Supplementary materials, methods, and data for "Proximity determines donor candidacy during DNA double-stranded break homology directed repair"

Submitted manuscript—all rights reserved

**The PDF file includes:**

Materials and Methods

Figs. S1 to S8

Table S2

References (38-55)

**Other Supplementary Materials for this manuscript include the following:**

Table S1. Tabulated data from Fig. 5 (provided as an Excel file)

**Materials and Methods**

Plasmid cloning and purification methods

Plasmids used in this study were generated using the NEBuilder HiFi DNA Assembly strategy (New England Biolabs, E2621L); assembly DNA fragments were derived using a combination of restriction digest, PCR amplification, or purchased (Integrated DNA Technologies, gBlocks).

Plasmids for lentivirus were raised in the *Stbl3* or *NEB Stable E. coli* strains. Plasmids used as donors in RaPID-seq genome editing HDR recombination experiments were raised in a *dam–/dcm– E. coli* strain (New England Biolabs, C2925H). All other plasmids were raised in *E. coli* strain *DH5α*. Plasmids were routinely purified from overnight *E. coli* cultures using conventional alkaline lysis with spin-column purification (QIAGEN, 27104; QIAGEN, 12941). Plasmid identity was validated using a combination of Sanger sequencing and Oxford Nanopore long-read sequencing (MicroSynth AG, Switzerland).

Purification of plasmids for RaPID recombination experiments

To purify plasmids for use as a donor in RaPID-seq HDR experiments, 100mL of overnight bacteria cultures (*dam–/dcm– E. coli*) were sedimented 10 minutes at 4°C at 4000rcf and the supernatant was discarded. Plasmids were isolated using conventional alkaline lysis with spin-column purification (QIAGEN, 12963) and eluted into low-EDTA TE buffer (10mM Tris-HCl, pH8; 0.1mM EDTA).

To further purify the plasmid from contaminating *E. coli* genomic DNA and *dam+* plasmids, the plasmids were digested overnight at 37°C by combining 100ng/µL plasmid, 0.2U/µL DpnI (New England Biolabs, R0176L), and 0.3U/µL BstEII (New England Biolabs, R3162M); the contaminants were then degraded by adding 0.2U/µL T5 Exonuclease (New England Biolabs, M0663L) and incubating for 2 hours at 37°C. The purified plasmid was isolated by combining the reaction mixture with QIAGEN Buffer BB, binding to a QIAGEN Plasmid *Plus* MidiPrep column, and washing per the manufacturer's protocol (QIAGEN, 12941). The purified plasmid was eluted in low-EDTA TE buffer.

To validate digestion of contaminating *E. coli* genomic DNA and *dam+* plasmids, a sample of each purified plasmid was digested with the (-*dam*)-specific restriction enzyme DpnII (New England Biolabs, R0543L) for 1 hour at 37°C. Samples of plasmid material from the initial isolation; DpnI, BstEII, T5 Exonuclease digest; and DpnII digest were analyzed by agarose gel electrophoresis stained with ethidium bromide to validate each processing step.

To quantify the purified plasmid, DpnII-digested plasmid was directly assayed using the Qubit dsDNA BR assay kit (ThermoFisher, Q32853). Purified plasmids were stored short-term at 4°C or long-term at -80°C in single-use aliquots.

General cell maintenance

The cells lines used in this study were human HEK293T (human embryonic kidney, SV40 large T antigen transformed) and human K562 (immortalized myelogenous leukemia). All cell cultures were maintained at 37°C and 5% $CO_2$ in a humidified incubator up to a maximum passage number P25. HEK293T cells were passaged every 2-4 days using Trypsin-EDTA

(ThermoFisher, 25200056) and seeded at a density of 2-5E4 live cells/cm2. K562 cells were maintained in log phase growth by passaging every 2-4 days, maintaining a cell density between 1E5 and 1E6 live cells/mL.

For HEK293T cells, culture media composition was: DMEM base media with GlutaMAX (ThermoFisher 10566016), 10 percent (v/v) standard fetal bovine serum (ThermoFisher, A5256701), and 100U/mL penicillin-streptomycin (ThermoFisher, 15140122). For wild-type K562 cells, culture media composition was: RPMI 1640 base media with GlutaMAX (ThermoFisher, 11875093), 10 percent (v/v) standard fetal bovine serum, and 100U/mL penicillin-streptomycin. For RaPID and Dam-only K562 cells, culture media composition was: RPMI 1640 base media, 10 percent (v/v) tetracycline-screened fetal bovine serum (ThermoFisher, A4736201), 100U/mL penicillin-streptomycin, and 1µg/mL puromycin (ThermoFisher, A1113803); for RaPID-BFP K562 cells, the RaPID/Dam media was further supplemented with 0.2µm sterile-filtered geneticin to 500µg/mL (ThermoFisher, 11811031).

Cell cultures were routinely tested for mycoplasma contamination using the MycoAlert Mycoplasma Detection Kit (Lonza, LT07-418). Cell line identity was verified using STR analysis (MicroSynth AG, Switzerland). Cell cultures were routinely cryopreserved at early passage (<P10) in a medium comprising 90 percent fetal bovine serum and 10 percent DMSO (v/v); components were individually 0.2µm sterile-filtered (Sarstedt, 83.1826.001; Sartorius, 17575-ACK) before aseptically combining.

### Preparation of lentivirus

Lentivirus was produced in HEK293T cells using conventional polyethylenimine (PEI) transfection of lentivirus packaging and transfer plasmids (*38*, *39*). Briefly, a transfection mixture was prepared by combining 18.7µg lentiviral packaging plasmid (Addgene #8455), 5.3µg envelope plasmid (Addgene #8454), and 16µg transfer plasmids with 120µg PEI in 1500µL OptiMEM media (ThermoFisher, 31985070). The transfection mixture was incubated 20 minutes at room temperature and thereafter added dropwise to HEK293T cells at 60 percent confluence in a 10cm dish. The lentivirus-enriched media from these cell cultures was harvested 24-, 48-, and 72-hours post-transfection; the harvested media was combined and 0.45µm filtered (Sarstedt, 83.1826) to remove residual cells and debris. The clarified lentivirus-enriched media was aliquoted directly or further concentrated using Lenti-X Concentrator (Takara, 631231) before aliquoting and storing at -80°C.

### Generation of stable-integration cell lines

K562 cells were transduced with lentivirus by limiting titration; 72 hours after transduction, the cells were treated with an appropriate selection antibiotic(s) to isolate single-integration polyclonal cell pools. Single-cells were sorted into 96-well plates and grown for 1-2 weeks before selecting surviving clones for expansion and further validation.

### CRISPR/Cas9 genome editing

CRISPR/Cas9 genome editing was performed as previously described using the Nucleofector X system (Lonza), 100µL cuvette format (*40*). Briefly, sgRNAs were first prepared by T7 *in vitro* transcription as previously described or synthetically produced (Synthego) (*28*). Thereafter, CRISPR/Cas9 and sgRNA ribonucleoprotein complexes (RNPs) were prepared by combining 200pmol Cas9-NLS, 240pmol sgRNA, and 4X Cas9 buffer (80mM, HEPES-KOH, pH 7.5; 600mM KCl; 4mM $MgCl_2$; 40% (v/v) glycerol). K562 cells (6-10E6 cells per Nucleofection) were sedimented and washed 3X with DPBS (ThermoFisher, 14190144). Thereafter, the CRISPR/Cas9 RNPs, cells, and Nucleofector Solution SF (Lonza, V4XC-2024) were combined in a cuvette and nucleofected using the FF-120 program code. After briefly incubating 5-10 minutes at room temperature, the cells were recovered into culture media and grown until harvesting.

### RAD51 proximity identification sequencing (RaPID-seq)

For RaPID-seq experiments, RaPID-K562 cells were treated for 20-24 hours before CRISPR/Cas9 genome editing with 80-100nM doxycycline (Tocris, 4090) at a density of 0.35E6 live cells/mL. The doxycycline treatment concentration was determined on a per-batch basis to ensure a %mCherry+ at genome editing of <20%; as previously reported, excessive Dam protein expression will cause non-specific genome-wide labelling and will substantially decrease DamID assay sensitivity (*11*). On the day of genome editing, a sample of cells was taken and stained with Hoechst (AnaSpec, 83218) before flow cytometry analysis to verify a normal dividing cell cycle distribution and *RaPID* gene expression (%mCherry+) levels.

To induce a targeted DSB in RaPID cells, CRISPR/Cas9 genome editing was performed as described above. In experiments targeting the *BFP* transgene, RNPs were prepared using 600pmol CRISPR/Cas9 protein and 720pmol sgRNA. After CRISPR/Cas9 genome editing, the cells were recovered into standard RaPID media supplemented to 1µM with Shield-1 (CliniSciences, AOB1848-5). For the RaPID-seq timecourse, cells were harvested at the indicated timepoints. Otherwise, the cells were harvested at 18h post-Nucleofection. For analysis of DSBR outcomes, 1E6 cells were seeded into standard RaPID media; the remaining cells were frozen at -80°C until extracting genomic DNA. For analysis of DSBR outcomes, the seeded cells were cultured for a further 2-3 days before harvesting.

### Genome-wide next generation sequencing (NGS) methods

All next-generation sequencing was performed in partnership with the Functional Genomics Center Zürich (FGCZ, Switzerland) and Genome Engineering and Measurement Lab (GEML, ETH Zürich). Paired-end sequencing was performed using a combination of the Illumina and Element Biosciences platforms. For DamID, the minimum number of reads was 10E6 reads per sample; for analysis of DSBR outcomes by targeted deep sequencing, the minimum number of reads was 1E5 reads per sample. A list of all DNA oligonucleotides and PCR primers is found in **Table S2**.

### Purification of genomic DNA

Genomic DNA was purified from cell pellets using the Puregene Kit (QIAGEN, 158043) or DNeasy Blood & Tissue Kit (QIAGEN, 69504) following the manufacturer's instructions; in all cases, the optional RNase A digestion step was performed as per the manufacturer's protocol. Purified DNA was eluted or rehydrated in low-EDTA TE buffer.

In case of the Puregene Kit, the genomic DNA was further fragmented to 30-50kbps by freeze-thawing 5X to reduce viscosity and improve pipetting precision. DNA was quantified using the Qubit dsDNA BR assay kit (ThermoFisher, Q32853) and stored at -20°C until use.

### Protein polyacrylamide gel electrophoresis (PAGE) and Western blot

Cells for protein extraction were harvested and rinsed 1X in DPBS (ThermoFisher, 14190144) before processing. Proteins were extracted by resuspending cell pellets in 1X RIPA buffer (Millipore, 20-188) supplemented with 1:100 Halt™ Protease Inhibitor Cocktail (ThermoFisher, 1861278) and incubating for 15-45 minutes on ice. Insoluble material was sedimented and discarded. Protein concentration was determined by the Bradford assay (VWR, M172-1L). Protein samples were standardized and denatured in 1X NuPAGE™ LDS Sample Buffer (ThermoFisher, NP0008) supplemented to 5 percent with 2-mercaptoethanol (Sigma-Aldrich, M6250) for 10 minutes at 70°C.

For protein PAGE, 42ug protein from each sample was loaded on a NuPage Bis-Tris™ gradient gel (ThermoFisher, NP0321BOX) and electrophoresed until protein ladder (ThermoFisher, 26623) was sufficiently separated. Proteins were transferred to a nitrocellulose membrane using the BioRad Trans-Blot® Turbo™ Transfer System. Membranes were blocked in 5 percent milk in TBST buffer (w/v). Proteins were visualized by incubating membranes with primary antibodies against V5-tag (Cell Signaling Technologies 13202S, Lot No. 5, diluted 1:1000) and GAPDH (Cell Signaling Technologies 97166S, Lot No. 5, diluted 1:5000) followed by secondary antibodies (LICOR 926-68072 and 926-32213; diluted 1:20000); all antibody dilutions were in 5 percent milk in TBST buffer (w/v). Blots were imaged using the LICOR Odyssey DLx Imager.

### DISCOVER-seq detection of genome editing target sites

Genome editing off-target detection by DISCOVER-seq were performed as previously described with minor modifications (*40*). Briefly, 20E6 K562 cells per sample were treated with CRISPR/Cas9-sgRNA RNPs by Nucleofection and formaldehyde-fixed after 12 hours. Chromatin immunoprecipitation was performed using an antibody against MRE11 (Novus Biologicals, NB100-142) followed by next generation sequencing. Target sites were identified with BLENDERv2 (https://github.com/cornlab/blender) using a MAPQ10 threshold followed by manual review of nominated genomic loci. For sgMulti1 analysis, DISCOVER-seq scores for sites within the same 25kbps bin were aggregated into a single site for comparison to RaPID-seq scores; the highest DISCOVER-seq score was chosen as the representative value.

### Strand-specific ChIP-seq (ssChIP-seq)

CRISPR/Cas9-treated cells were harvested as previously described and stored at -80°C until processing. Chromatin immunoprecipitation was performed as previously described using antibodies targeting V5-tag (Abcam ab9116, Lot No. GR3361888-2) or RAD51 (Novus

Biologicals NB100-148, Lot No. 44265) (*40*). Strand-specific libraries for next-generation sequencing were prepared from the purified ChIP material using the xGen ssDNA & Low-Input DNA Library Prep Kit (Integrated DNA Technologies, 10009859) according to the manufacturer instructions.

DNA methyltransferase identification sequencing (DamID-seq)

Dam-labelled dsDNA GATC fragments (N6-Methyladenosine, Gm6ATC) were enriched from genomic DNA using the previously reported DamID methods with minimal modifications; the DamID adapter was prepared by annealing DNA oligomers (DamID-AdR_fwd; DamID-AdR_rev) purchased from Integrated DNA Technologies (*7*, *8*).

Briefly, purified genomic DNA was combined with *+dam* Lambda DNA (New England Biolabs, N3011L) spike-in at a ratio of 8pg or 40pg per 100ng genomic DNA. The DNA was then treated with shrimp alkaline phosphatase (New England Biolabs, M0371L) for 1 hour at 37°C to prevent enrichment of non-biological DNA breaks resulting from DNA purification and/or sample freezing. Thereafter, the DNA was then treated with DpnI (New England Biolabs, R0176L) for 10 hours at 37°C to expose sites of Dam-labelled DNA to which the DamID adapter was ligated overnight (New England Biolabs, M0202L) at 16°C. A final digestion with DpnII (New England Biolabs, R0543L) was performed for 1h at 37°C to prevent PCR enrichment of non-Gm6ATC sites. This DamID-digested DNA was immediately used in DamID-PCR enrichment or stored at -20°C.

To enrich Dam-labelled dsDNA fragments, 40µL PCR reactions were prepared by combining: 8µL DamID-digested DNA, 20 µL MyTaq Mix 2X (Bioline, BIO-25041), 1.25µM DamID-PCR primer, and water. The PCR amplification was performed for a total of 22 cycles as follows: initial extension: 10min at 68°C; amplification group 1 (1 cycle): 1min at 94°C, 5min at 65°C, 15min at 68°C; amplification group 2 (4 cycles): 1min at 94°C, 1min at 65°C, 10min at 68°C; amplification group 3 (17 cycles): 1min at 94°C, 1min at 65°C, 2min at 68°C; final extension: 5min at 68°C. Thereafter, 5 µL of each PCR reaction was analyzed by agarose gel electrophoresis for a characteristic DamID smear of 200-2000bps.

The remaining DamID-PCR product was purified by combining with 1.5X ratio (v/v) solid-phase reversible immobilization (SPRI) beads (Cytiva 29343052), washing with 85 percent (v/v) ethanol/water solution, and eluting in low-EDTA TE buffer. Purified DamID-PCR product was quantified using the Qubit dsDNA BR assay kit (ThermoFisher, Q32853); up to 1000ng of purified DamID-PCR product was sheared to a mean fragment size of 300-600bps using the Covaris LE220-plus focused-ultrasonicator system. The sonicated material was directly used in preparing Illumina next-generation sequencing libraries using the NEBNext Ultra II workflow (New England Biolabs, E7645L) following the manufacturer's instructions.

Pre-PCR for targeted deep sequencing to determine DSBR outcomes (amplicon-NGS)

To assess DSBR outcomes, targeted amplicon-NGS of DSBs was performed as previously described (*13*). Briefly, a stubbed-amplicon product was generated by an initial PCR amplification and purified before being used in a subsequent NGS-indexing PCR.

For the direct stubbing PCR (*HBB* locus), 50µL reactions were prepared by combining: 100ng purified genomic DNA, 25µL NEBNext Ultra II Q5 Master Mix (New England Biolabs No. M0544L), 500nM each forward and reverse primer, and water. The stubbing PCR protocol was as follows: initial denaturation: 3min at 98°C; amplification (22 cycles): 10sec at 98°C, 30sec at 64°C, 40sec at 72°C; final extension: 2min at 72°C. Thereafter, the stubbing PCR reactions were purified using SPRI beads.

Where necessary, a nested PCR strategy was used wherein an approximately 1kbps region was PCR amplified, purified, and then used as a template for a subsequent stubbing-PCR amplification. The nested-PCR reactions were purified using SPRI beads (ratio 1.0X). The subsequent stubbing-PCR was amplified for only 3 cycles but otherwise prepared as previously discussed.

For genomic loci *OR7E39P* and *OR7E89P*, the nested-PCR was prepared as follows: 100ng purified genomic DNA, 25µL AmpliTaq Gold 360 (ThermoFisher 4398881), 500nM each forward and reverse primer, and water. The nested-PCR protocol was as follows: initial denaturation: 10min at 95°C; amplification (22 cycles): 30sec at 95°C, 2min at 56°C (*OR7E89P*) or 60°C (*OR7E39P*), 1min 30sec at 72°C; final extension: 7min at 72°C.

For genomic loci *HBB* the nested-PCR strategy was applied in RaPID-seq plasmid experiments to specifically enrich genomic DNA sequences and exclude plasmid DNA sequences. The nested-PCR was prepared as follows: 200ng purified genomic DNA, 25µL NEBNext Ultra II Q5 Master Mix (New England Biolabs No. M0544L), 500nM each forward and reverse primer, and water. The nested-PCR protocol was as follows: initial denaturation: 3min at 98°C; amplification (22 cycles): 10sec at 98°C, 30min at 64°C, 40sec at 72°C; final extension: 2min at 72°C.

<u>Indexing PCR for targeted deep sequencing to determine DSBR outcomes</u>

For NGS-indexing PCR, 50µL reactions were prepared by combining: 15µL purified stubbing-PCR product, 25µL NEBNext Ultra II Q5 PCR Master Mix, 1µM each forward and reverse primer (New England Biolabs E7600S), and water. The indexing PCR protocol was as follows: initial denaturation: 30sec at 98°C; amplification (3-5 cycles): 10sec at 98°C, 1min 15sec at 65°C; final extension: 5min at 65°C. Thereafter, the stubbing PCR reactions were purified using SPRI beads and pooled for next-generation sequencing.

<u>Polyclonal donor pool generation for one-to-all genome-wide mapping of endogenous HDR pairs</u>

To generate genome-wide HDR-donor (*+GFP*) polyclonal pools for endogenous HDR experiments, RaPID-BFP cells were transduced at a MOI <0.15 with a lentivirus containing an expressed mCarmine gene and a non-expressed GFP donor. Cells were cultured for at least 4 days on RaPID-BFP media before isolating +mCarmine/-GFP cells by FACS using the Sony SH800 cell sorter system. These RaPID-BFP cell pools with polyclonal single-GFP donor integrations (RaPID-BFP+GFP cells) were further expanded and cryopreserved before performing further experiments.

Unbiased one-to-all endogenous HDR recombination loci pair screening

RaPID-BFP+GFP cell pools were cultured on standard RaPID media without geneticin. *BFP* was targeted in 10E6 RaPID-BFP+GFP cells using the previously described using CRISPR/Cas9 genome editing. The edited cells were cultured for at least 7 days before isolating GFP+ cells by FACS using the BD FACSAria™ III Cell Sorter (ETH Flow Cytometry Core Facility). Sorted cells were allowed to recover in culture before cryopreserving and harvesting for analysis.

Library preparation for mapping of lentivirus insertion sites (LentiMap)

Based on the GUIDE-seq enrichment strategy, we designed a protocol to map lentiviral integration sites (*20*). Briefly, 1µg of genomic DNA was fragmented to 500-700bp using the Covaris LE220-plus focused-ultrasonicator system, followed by end-repair, and ligation to the P5 adapter—having an 8bp UMI sequence and an 8bp sequencing barcode—using the NEBNext Ultra II workflow (New England Biolabs, E7645L) according to the manufacturer's instructions.

Next, PCR enrichment of lentivirus insertion sites was performed using Q5 Hot Start High-Fidelity DNA Polymerase (New England Biolabs, M0493L) for 14 cycles with PCR primer LentiMap_amp, followed by SPRI bead purification (ratio 0.80X). A second PCR was performed for 14 cycles to complete the P7 Illumina TruSeq adapter and attach the second sequencing barcode. Due to high background amplification from the P5 primers, library concentration was estimated by qPCR. Libraries were sequenced using the NextSeq 2000 (Illumina) platform with the following strategy: Read 1: 50bp; Index1: 8bp; Index2: 16bp; Read2: 150bp; library loading concentration: 300pM; PhiX 10-20 percent. The samples were demultiplexed, keeping the UMI information in a third read for subsequent data analysis.

Analysis of post-DSBR amplicon-NGS

Targeted deep sequencing paired-end FASTQ data were processed using the *CRISPResso2* (version 2.3.1) software to summarize DSBR event species (*41*). HDR outcomes were manually annotated by identifying DSBR outcomes with gene conversion tracts corresponding to donor DNA sequences.

Analysis of RaPID-seq for homology search interactions (analyzeRaPID)

FASTQ datafiles from RaPID-seq Illumina NGS were preprocessed using three consecutive rounds of *cutadapt* (version 4.6) to remove Illumina TruSeq adapters, DamID adapters, and known amplicon-contaminants (*42*). The preprocessed reads were then aligned to the human genome (*GRCh38*) and Lambda phage genome using *bowtie2* (version 2.5.2) and converted to bam format using *samtools* (version 1.19) (*43–45*). Counts of reads aligning to Lambda phage genome were extracted for spike-in correction. Reads aligning to *GRCh38* were counted into DpnI restriction digest spans (GATC windows) using *htseq-count* (version 2.0.4) (*46*).

In RaPID-seq plasmid experiments, FASTQ datafiles after *cutadapt* processing were first aligned to the plasmid sequences using *bowtie2* and converted to bam format using *samtools*. Reads not aligning to the plasmid were extracted and then aligned to *GRCh38* and processing proceeded as previously indicated.

Preprocessed data were aggregated using our analyzeRaPID (https://github.com/yehcd/rapid-tools) software to perform spike-in correction and background subtraction; key dependencies include *GenomicRanges* (version 1.52.1), *rtracklayer* (version 1.60.1), and *tidyr* (version 1.3.0) (*47–49*). For each experimental series, the non-targeting CRISPR/Cas9 RNP sample was used as the common background sample. Briefly, Lambda spike-in counts were used to adjust the GATC window scores from *htseq-count* processing. Thereafter, scores from +DSB and -DSB samples were subtracted and normalized based on read depth and GATC window size (FPKM, fragments per kilobase per million-mapped reads), yielding the RaPID FPKM score. To aggregate RaPID-seq values from several independent experiment series, values were rescaled by global maximum to ±1 range. Outputs of analyzeRaPID processing were binned to 25kbps or 100kbps windows and exported as bigWig datafiles, which were plotted using *ggplot2* (version 3.5.1) (*50*).

### Analysis of strand-specific ChIP-seq (ssChIP-seq)

FASTQ datafiles from Illumina NGS were preprocessed using two consecutive rounds of *cutadapt* to remove Illumina TruSeq adapters and known amplicon-contaminants (*42*). The preprocessed reads were then aligned to the human genome (*GRCh38*) using *bowtie2* and converted to bam format using *samtools* (*43–45*). Normalized coverage was calculated as counts per million (CPM) using *deepTools* (*bamCoverage*, version 3.5.4) and plotted using *ggplot2* (*50*, *51*).

### Analysis of sequence homology

Sequence homology analysis was performed using EMBOSS *clustalo* (version 1.2.4) (*52*, *53*). Alignment and percent identity scores were extracted for further analysis.

### Analysis of HiC chromatin conformation and RaPID-seq cross comparison

HiC contact matrices for wild-type K562 cells were retrieved from the ENCODE project (ENCFF621AIY), and individual virtual-4C views with balanced normalization corresponding to CRISPR/Cas9 DSB sites were extracted using *strawr* (version 0.0.92) (*16*, *54*).

A region corresponding to DSB±150kbps was masked to exclude effects resulting from DSBR end processing. Using the HiC expected interaction frequency model, the RaPID-seq and observed HiC values were rescaled such that the expectation value at the DSB±150kbps position is 1.0. For observed HiC values, this was accomplished by direct rescaling to the target value; for RaPID-seq, this was accomplished by first generating a naïve distance-proximity expected interaction score for each RaPID-seq GATC window using the HiC expected interaction model followed by direct rescaling to the target value.

Analysis of LentiMap

FASTQ datafiles from RaPID-seq Illumina NGS were first annotated with UMI barcodes using *umitools* (version 1.1.4). Thereafter, *cutadapt* was used to remove Illumina TruSeq adapters and extract lentivirus-specific reads, which were then aligned to *GRCh38* and deduplicated using *bowtie2*, *samtools*, and *umitools* (*42–44*, *55*). Normalized coverage was calculated using *deepTools* (*bamCoverage*, version 3.5.4) as counts per million (CPM) in 25kbps bins and plotted using *ggplot2* (*50*, *51*).

To identify *GFP* donor sites, a threshold was applied to the resulting LentiMap coverage datafile. Overrepresented *GFP* donor sites in the untreated *BFP+GFP* pool were identified by LentiMap analysis; sites in the top 2 percentile (LentiMap CPM ≥ 20) were excluded from further analysis as overrepresented or non-specific amplification. This same threshold value was used to identify sites of *bona fide* recombination between the targeted *BFP* gene and *GFP* donor sequences in CRISPR/Cas9 RNP-treated samples. For background comparison, *GFP* donor sites in CRIPSR/Cas9-treated conditions with a LentiMap CPM ≥ 15 were also plotted but not included in Spearman's correlation (ρ) analysis.

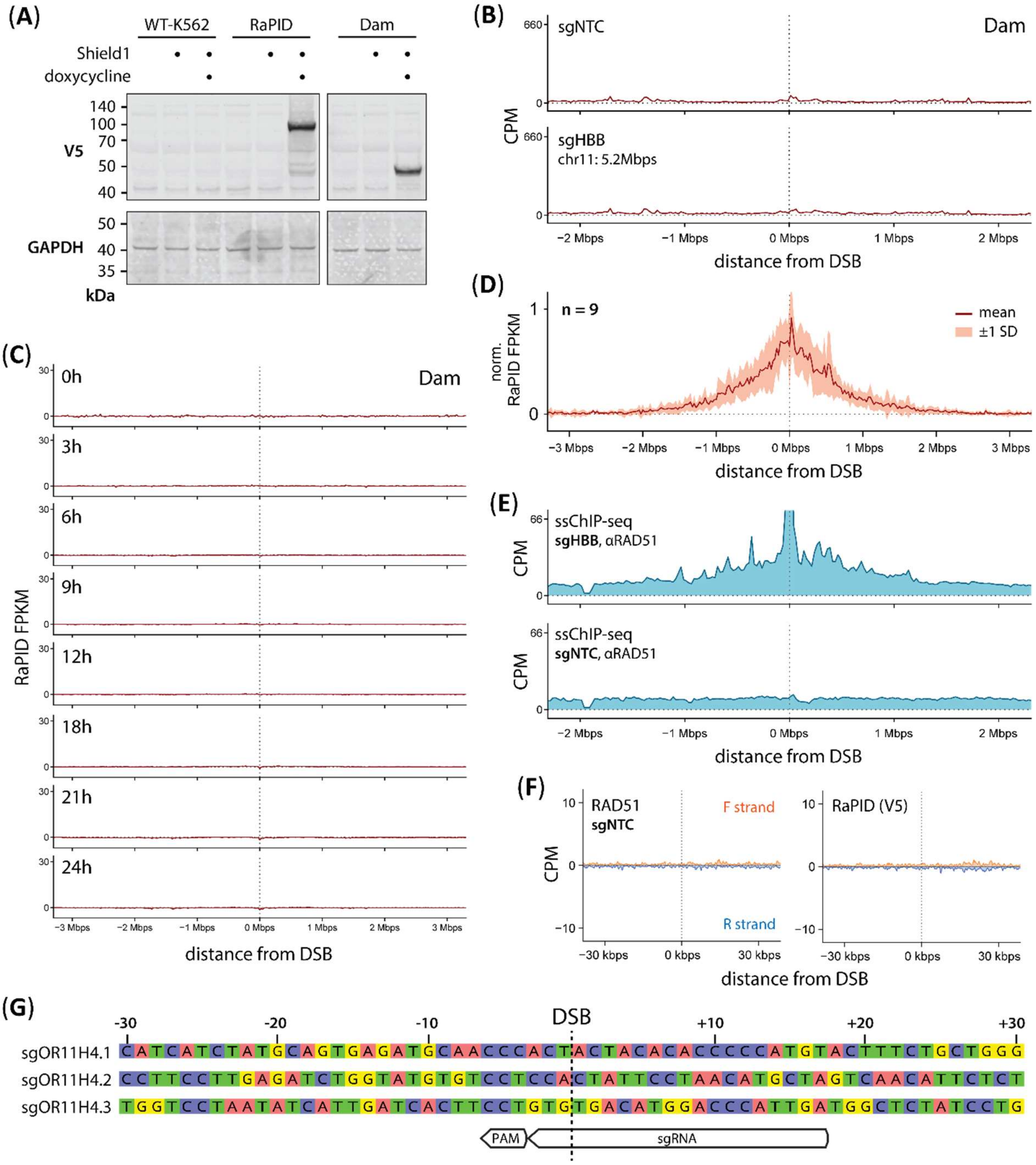


**Fig. S1.**

**RaPID-seq signal is highly specific to DSB formation and Dam fusion to RAD51 protein. (a)** Western blot for RaPID and Dam-only K562 cell lines demonstrates detectible protein levels only in the presence of both doxycycline and Shield1 treatment. **(b)** DamID on Dam-only K562 cells at 18h post-Nucleofection showed no signal irrespective of DSB presence. **(c)** Timecourse of DamID signal for a DSB at *HBB* in Dam-only K562 cells. **(d)** Average of normalized RaPID-seq homology search profiles for the CRISPR/Cas9 *sgMulti1* at the 9 unique target sites that with sufficient linear distance separation (>3Mbps); band is ±1SD. **(e)** Y-axis zoom of RAD51 ssChIP-NGS with (sgHBB) or without (sgNTC) CRISPR/Cas9 targeting a DSB at *HBB*. **(f)** Strand-specific analysis of ssChIP for RAD51 and RaPID (V5) in the absence of a DSB, neither RAD51 nor RaPID are recruited to the *HBB* locus. **(g)** Comparison of genomic sequence immediately around the 3 different CRISPR/Cas9 sgRNAs targeting *OR11H4*.

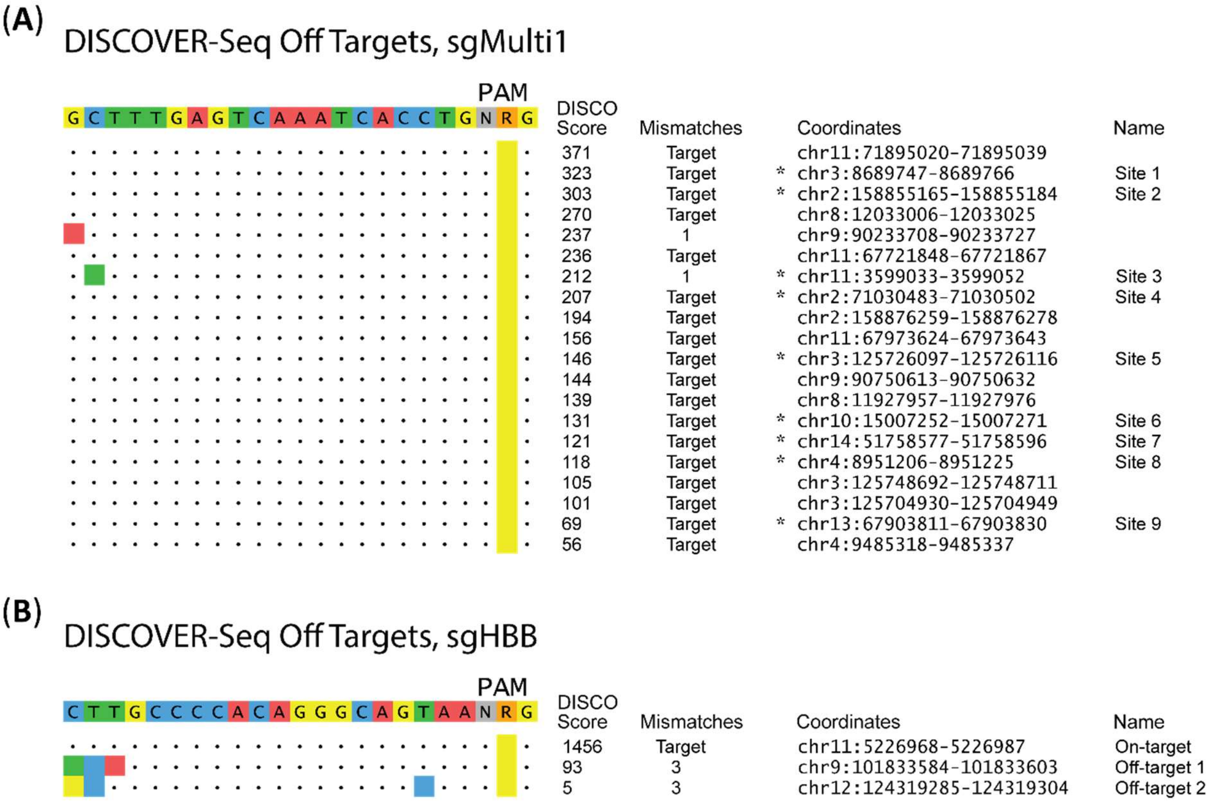


**Fig. S2.**

**DISCOVER-seq and BLENDERv2 analysis of genome editing target sites**. K562 cells edited with CRISPR/Cas9 RNPs using guide RNAs **(a)** *sgMulti1* and **(b)** sg*HBB*. Only sufficiently separated sites (*) that have non-overlapping RaPID-seq profiles were used in the analyses of **Fig. 1g and S1g**.

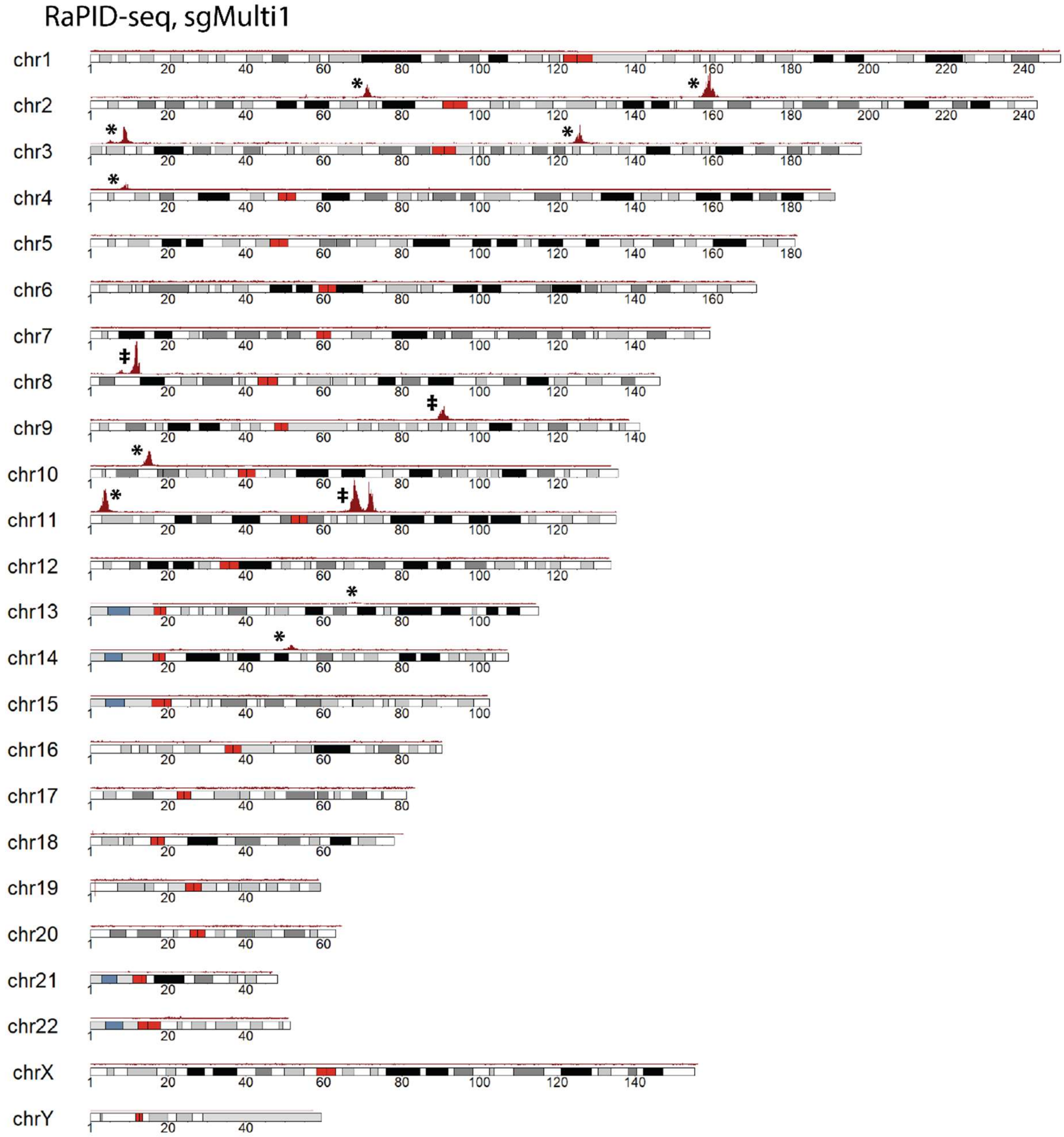


**Fig. S3.**

**Genome-wide view of homology search profiles for the CRISPR/Cas9 targeting with *sgMulti1*.** Selected DSB target sites (*) were identified using DISCOVER-seq (**Fig. S2**). Regions (‡) with clusters of high efficiency CRISPR/Cas9 target sites as detected by DISCOVER-seq also have high RaPID-seq search signal but were excluded from analysis due to overlapping homology search profiles.

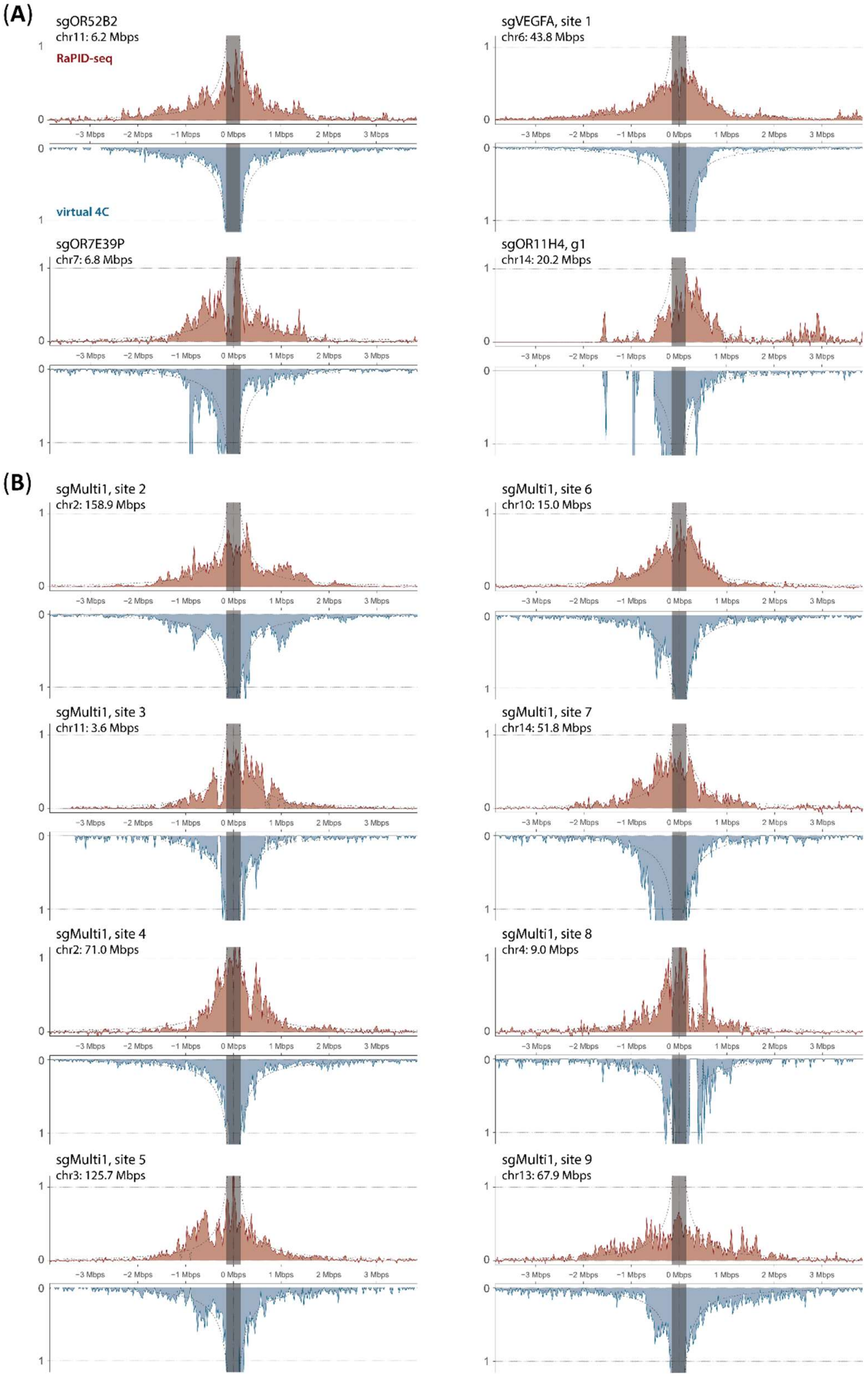


**Fig. S4.**

**Comparison between unperturbed K562 cell chromatin conformation (HiC) and DSBR homology search (RaPID-seq) profiles.** Dotted line curve: expected search profile for a naïve linear distance model of DNA interaction; masked region is DSB±150kbps. Analysis of **(a)** additional CRISPR/Cas9 target sites and **(b)** 8 other *sgMulti1* target sites identified by DISCOVER-seq.

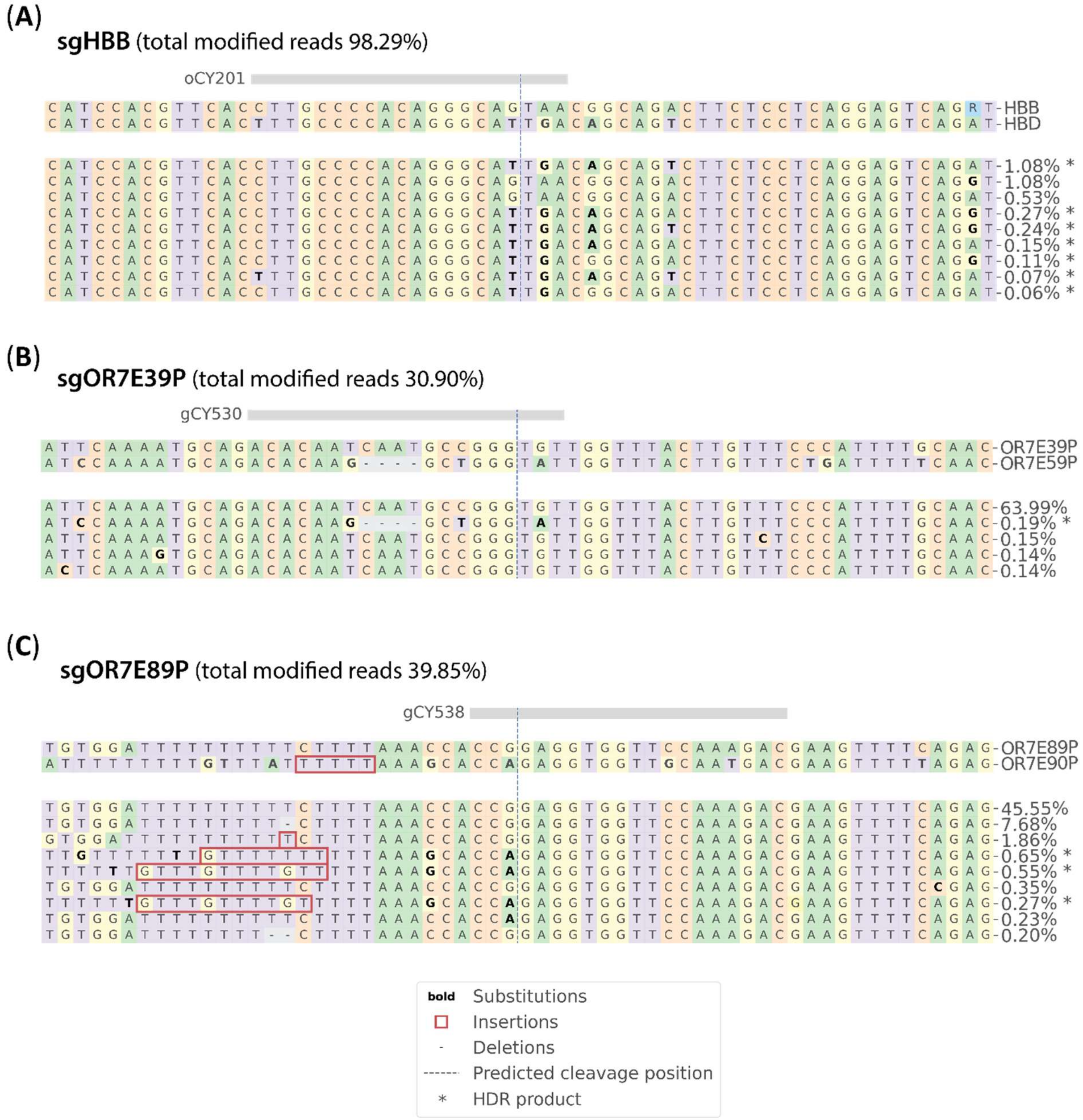


**Fig. S5.**

***CRISPResso2* analysis of DSBR outcomes.** Exemplar HDR products (*) for DSBs targeted at **(a)** *HBB*, **(b)** *OR7E39P*, and **(c)** *OR7E89P*. The endogenous HDR donor gene is indicated for each targeted site; being *HBD*, *OR7E59P*, and *OR7E90P*, respectively. Percent frequencies given as fraction of all reads, including unmodified alleles.

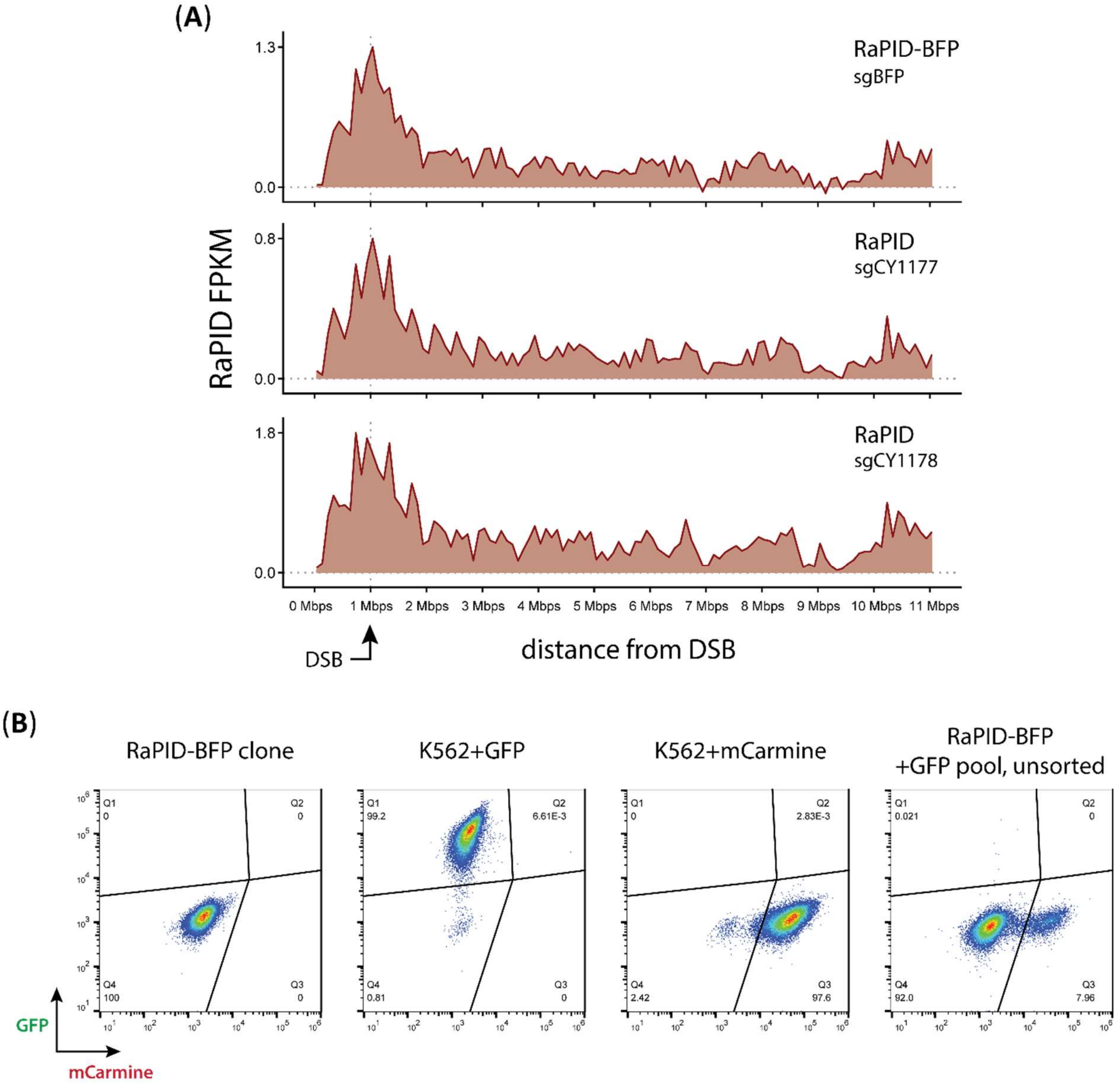


**Fig. S6.**

**Extended data for unbiased genome-wide *in situ*/*in vivo* one-to-all HDR recombination analysis by *BFP* to *GFP* gene conversion. (a)** Comparison of homology search profiles at the monoallelic *BFP* in RaPID-BFP cells and the same locus in the parental RaPID cells. Note that in the parental RaPID cells, no *BFP* gene exists. **(b)** Estimation of lentivirus transduction MOI for the +GFP donor pool. RaPID-BFP (6E6 cells) were transduced with lentivirus bearing a non-expressed *GFP* donor gene and an expressed *mCarmine* gene; %mCarmine+ corresponds to MOI.

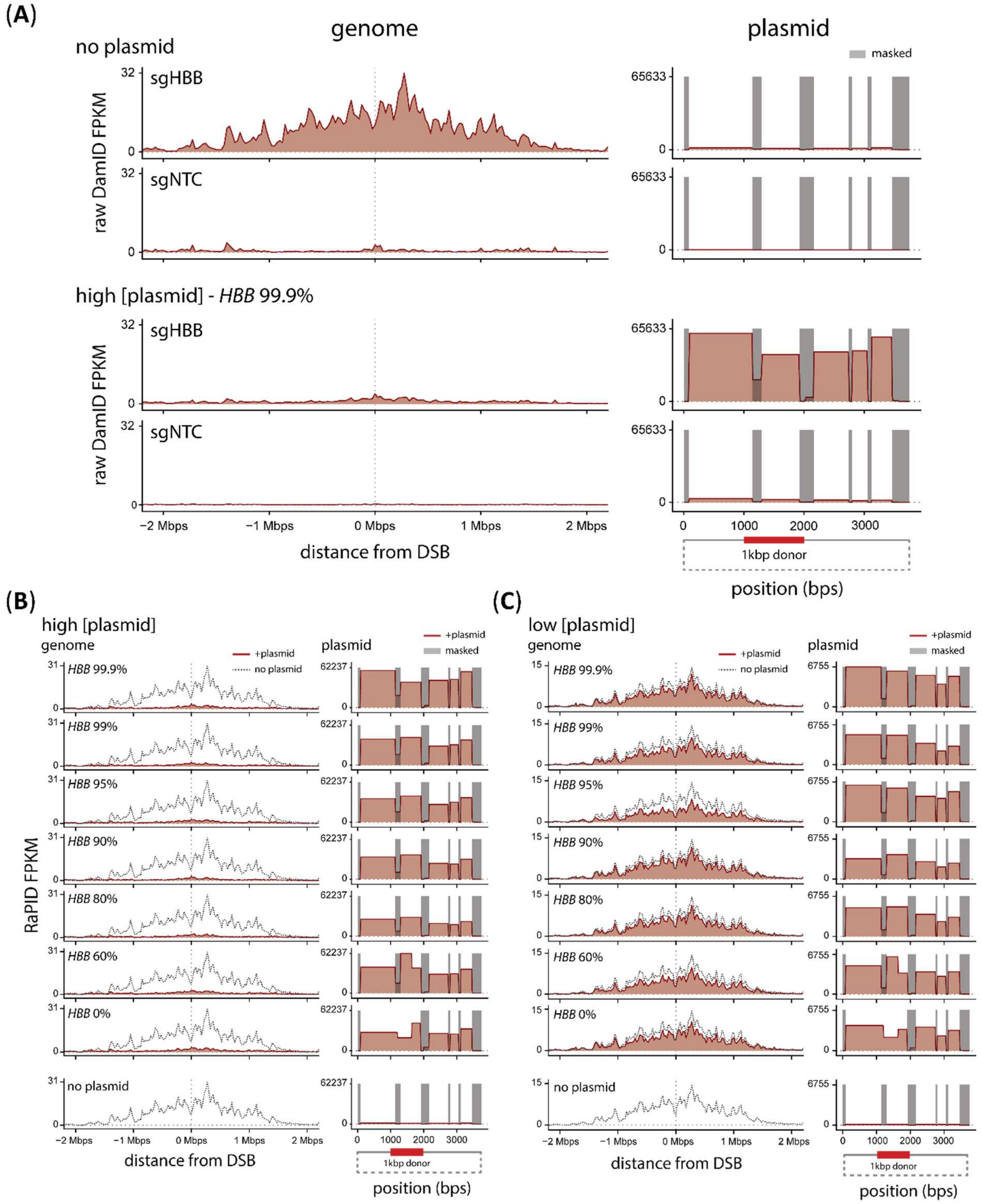


**Fig. S7.**

**Extended data for RaPID-seq for a DSB at *HBB* in cells treated with plasmid donors. (a)** RaPID-seq data before final background subtraction in cells treated with ±targeting CRISPR-Cas9 RNP at *HBB* and treated with ±plasmid donor (99.9% homology to *HBB*). No plasmid condition data was aligned to the *HBB 99.9%* plasmid donor to determine background levels. **(b)** RaPID-seq for a DSB at *HBB* in cells treated with 100ng/µL (high) or **(c)** 12.5ng/µL (low) concentrations of exogenous donor plasmids with varying *HBB* homology. Gray dotted line indicates homology search profile in absence of exogenous plasmid; non-amplifiable regions (<200bps) on plasmid are masked from analysis.

Submitted manuscript—all rights reserved

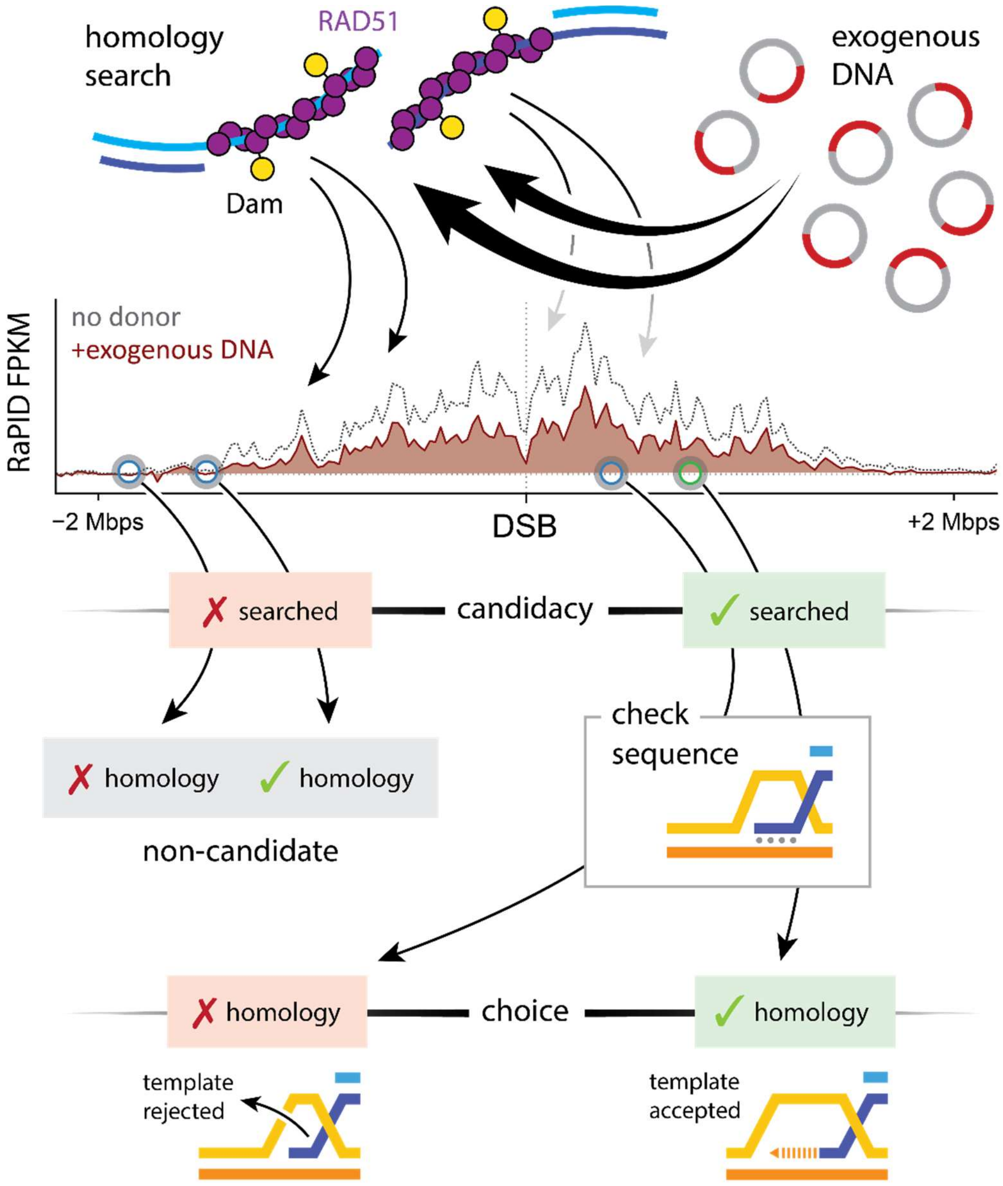


**Fig. S8.**

**Model for competition between endogenous and exogenous DNA during DSBR homology search.** Search predominantly probes the DSB local region and is further constrained by chromatin conformation. Exogenous plasmid DNAs are abundant and highly accessible, thereby competing with endogenous search in a concentration-dependent and homology-independent manner. Successful recombination between a DSB and a template first requires candidacy, which is dictated by proximity to the break site. DNA sequences that are not searched cannot serve as donor templates. Only those searched templates that pass a further DSB-template sequence comparison step unlock recombination where information is copied from a donor template into the break site.

| name | aliases | sequence (5' to 3') |
|---|---|---|
| DamID-AdR_fwd | oCY182 | CTAATACGACTCACTATAGGGCAGCGTGGTCGCGGCCGAGGA |
| DamID-AdR_rev | oCY183 | TCCTCGGCCG |
| DamID-PCR | oCY184 | GGTCGCGGCCGAGGATC |
| HBB_ngs_F | oCY856 | CTTTCCCTACACGACGCTCTTCCGATCTACATGCCCAGTTTCTATTGGT*C |
| HBB_ngs_R | oCY857 | GGAGTTCAGACGTGTGCTCTTCCGATCTGGGTTGGCCAATCTACTC*C |
| HBB_plasmidPreamp_F | oCY1184 | ATGCTTAGAACCGAGGTAGAGTT*T |
| HBB_plasmidPreamp_R | oCY1188 | GTAGAGTTTTCATCCATTCTGTC*C |
| HBB_plasmidNgs_F | oCY1186 | CTTTCCCTACACGACGCTCTTCCGATCTAGGGTTGCCCATAACAG*C |
| HBB_plasmidNgs_R | oCY1187 | GGAGTTCAGACGTGTGCTCTTCCGATCTGGCTGAGGGTTTGAAGT*C |
| OR7E39P_preamp_F | oCY549 | GGTGACAACCGCGTACAT |
| OR7E39P_preamp_R | oCY551 | CGGTCTCAGCTGTTCTACAGTGTT |
| OR7E39P_ngs_F | oCY593 | CTTTCCCTACACGACGCTCTTCCGATCTTCAGAACTGCAAAATATTGCT |
| OR7E39P_ngs_R | oCY594 | GGAGTTCAGACGTGTGCTCTTCCGATCTGGCTCTGCATAGTTTGG |
| OR7E89P_preamp_F | oCY569 | TATGCCTTTAACTGGAATGGGTC |
| OR7E89P_preamp_R | oCY570 | ATTTGTATGCTGTTGATGGACAAC |
| OR7E89P_ngs_F | oCY587 | CTTTCCCTACACGACGCTCTTCCGATCTGGGAGCTTACAAATGGTG |
| OR7E89P_ngs_R | oCY588 | GGAGTTCAGACGTGTGCTCTTCCGATCTTCACAATTAATTTCTGTTCTTACTC |
| LentiMap_amp | oCY1189 | GGAGTTCAGACGTGTGCTCTTCCGATCTATCTGCTTTTTGCTTGTACTG*G |

**Table S2.**

**DNA oligonucleotides used in this study.** Description of sequences used for PCR amplification of DamID libraries, targeted amplicon-NGS (*HBB*, *OR7E39P*, *OR7E89P*), and LentiMap. Phosphorothioate bond indicated by *.
